# Crosstalk between Echinoid and Sidekick, two IgCAM proteins, modulates Adherens Junction dynamics and tissue remodelling

**DOI:** 10.64898/2026.04.22.720164

**Authors:** Fleur Chelemen, Maria Lluisa Espinàs, Marta Llimargas

**Affiliations:** Department of Cells and Tissues Institut de Biologia Molecular de Barcelona, IBMB-CSIC. Parc Científic de Barcelona Baldiri Reixac, 10-12. 08028 Barcelona, Spain; Institute for Research in Biomedicine (IRB Barcelona), The Barcelona Institute of Science and Technology (BIST), Baldiri Reixac 10, 08028, Barcelona, Spain

## Abstract

Adherens junctions (AJs) undergo dynamic remodelling during epithelial morphogenesis, requiring precise coordination between adhesive proteins, intracellular adaptors, and cytoskeletal regulators. In addition to cadherins, which mediate core cell-cell adhesion and connect junctions to the actin cytoskeleton, other adhesion molecules from the immunoglobulin superfamily (IgCAM) also contribute to AJ organisation. In *Drosophila*, Echinoid (Ed), a nectin-like IgCAM, localises along the entire AJ, whereas Sidekick (Sdk), another IgCAM, is predominantly enriched at tricellular adherens junctions (tAJs). Although both proteins interact with overlapping intracellular partners, how they functionally relate to one another has remained unclear. Here, we investigate the spatial, molecular, and functional interplay between Sdk and Ed during embryonic epithelial morphogenesis. Using SRRF-imaging we show that Sdk and Ed frequently colocalise at tAJs but also display adjacent or spatially separated distributions; together with proximity-labelling experiments, these results suggest that Sdk and Ed engage in transient and dynamic associations rather than forming a stable complex. Functional analyses reveal that they influence each other’s accumulation, indicating bidirectional regulatory interactions. We find that Sdk modulates Ed levels along the entire AJs and affects Ed enrichment at tAJs. We provide evidence that this regulation involves changes in Ed intracellular trafficking, suggesting that Sdk modulates Ed levels at AJs at least in part by controlling its trafficking. Genetic analyses uncover previously unreported contributions of *ed* to tracheal cell intercalation and of *sdk* to dorsal closure, and reveal strong genetic interactions between the two genes, indicating cooperative yet context-dependent functions. Consistent with this, we find that Sdk and Ed converge on shared intracellular adaptor proteins, including Canoe and Polychaetoid, modulating their levels and junctional enrichment. Together, our findings support a model in which dynamic, multi-component protein complexes assemble at bicellular and tricellular AJs, integrating shared and junction-enriched components that engage in multiple, simultaneous, and mutually influencing interactions. This interconnected network would confer the spatiotemporal robustness and flexibility required to support the distinct cellular behaviours underlying tissue-specific remodelling.

## Introduction

Epithelial tissues are present in all organisms where they perform vital functions including protection, separation, secretion, absorption, filtration or transport. Cells forming epithelia are tightly attached to one another and exhibit marked apicobasal polarity. Adhesion and polarity are mediated by distinct protein complexes that localise to specific types of junctions. A principal component of epithelial Adherens Junctions (AJ) is E-Cadherin (Ecad), a calcium-dependent homophilic adhesion protein that links to the actin cytoskeleton, regulating epithelial organisation, cell rearrangements and cell shape changes (1–4).

In flat epithelia, bicellular AJs (bAJs) connect pairs of neighbouring cells, while tricellular AJs (tAJs) form at points where three of more cells meet, resulting in distinct architectural organisations. These architectural differences both confer and demand functional specificity, with tAJs acting as points of high tension as they need to withstand considerable mechanical forces (5–9). bAJs and tAJs also differ in their molecular composition: several proteins are uniformly present along the entire AJ, whereas others are enriched at tricellular AJs (5). Among them, Sidekick (Sdk) stands out as the only protein identified so far with a predominantly specific localisation at tAJs in most epithelial tissues analysed, including the trachea, epidermis, or follicular epithelia. Sdk encodes a member of the immunoglobulin cell-adhesion molecule (IgCAM) superfamily with a large extracellular domain containing 6 immunoglobulin (Ig) domains and 13 fibronectin type III (FNIII) domains, and an intracellular domain containing a C-terminal PDZ-binding domain. Sdk enrichment at tAJs is modulated by different, likely complementary, mechanisms, including the contributions of particular protein regions (10, 11), and also by tension (12). Nevertheless, Sdk is also detected in bAJs in a few epithelial contexts, such as the epidermis at germ band extension and the salivary glands at larval stages (12, 13), indicating its ability to localise along the entire AJs.

AJs are force-sensor structures and provide cell-cell adhesion through homotypic interactions between the Ecad molecules. However, besides Cadherins, other adhesion molecules are also present at AJs, which also contribute to AJs functions (5). For example, Sdk plays a role in resolving cell rearrangements events in various epithelial tissues during morphogenesis (12–14). AJs contain a second IgCAM family member from the Nectin subfamily, called Echinoid (Ed) in *Drosophila*, which localises along the entire bAJs and tAJs. Ed plays key roles in several developmental events including dorsal closure, head involution, neurogenesis or eye formation, and its differential accumulation between contacting cells induces the formation of contractile actomyosin cables at interfaces, which are required for proper morphogenesis (15–17). Similar to Sdk, Ed contains 7 Ig domains and one FNIII domain in the extracellular region and a PDZ-binding domain in its intracellular C-t region (18, 19). The extracellular domain of Ed can engage in both homophilic and heterophilic interactions (20), allowing it to exert its activity in coordination with other adhesion molecules.

We and others proposed that Sdk’s activity is mediated through interactions with various intracellular factors that modulate the actomyosin cytoskeleton, thereby regulating tissue remodelling. Indeed, actin-binding and actin-regulating proteins such as Canoe (Cno), Polychaetoid (Pyd), WAVE complex and MRLC have been identified by different approaches as interacting with Sdk (10, 12, 14, 21). Similarly, the intracellular PDZ-binding domain of Ed has been shown to interact with different proteins, including Bazooka and Cno (19), the latter of which also interacts with Sdk (10, 12).

Considering the molecular similarities between Sdk and Ed and their ability to interact with common partners, we asked how their differential enrichment in bAJs and tAJs regulates AJs activities in different tissues, and we explored potential interactions between them. We first observed that Ed is also slightly enriched at tAJs, where it tends to colocalise with Sdk. Our data suggest that the two proteins can form a complex, although this may be a transient one. Our findings suggest that each IgCAM can influence the other’s distribution between bAJs and tAJs. We find that Sdk affects Ed intracellular trafficking, suggesting that Sdk may regulate Ed accumulation along AJs through this mechanism. Our analysis reveals previously unreported minor roles for *ed* in tracheal intercalation and for *sdk* in dorsal closure, processes associated with *sdk* and *ed*, respectively. In addition, the phenotypes of *sdk; ed* double mutants suggest that both genes act cooperatively during the morphogenetic processes of tracheal intercalation and dorsal closure. Consistent with this, we find that reducing *sdk* and *ed* activity leads to a significant reduction in the tAJ enrichment of Pyd and Cno. Additionally, we identify a regulatory relationship between *ed* and *pyd.* Altogether, our findings support a model in which local proportions of Sdk and Ed dynamically modulate the molecular composition of AJs to achieve the specific cellular behaviours required for tissue remodelling.

## Results and discussion

### 1. Analysis of Sdk and Ed localisation

Sdk and Ed have been shown to localise at AJs, with Sdk being predominantly restricted to tAJs in most epithelia analysed (12–14, 22) and Ed localising in a broader pattern including both bAJs and tAJs (19). To investigate their relative localisation, we performed immunostainings in embryos using an antibody against Sdk and the pattern of GFP provided by a knock-in *ed*-tagged allele (ed^MI01552-GFSTF.1^, (23), from now on *ed^GFP^*). We analysed two different tissues, the epidermis and the trachea, at stage 14.

Conventional confocal microscopy indicated that the two proteins largely colocalise in a single spot at tAJs (Fig 1A). To more precisely determine and quantify their relative localisation we employed Super-resolution radial fluctuation (SRRF) imaging (24) (Fig 1B). This analysis uncovered a more complex spatial relationship between Sdk and Ed than initially expected. We classified the relative localisation of the two proteins at tAJs into three main categories based on the distance between the center of the fluorescence for each protein in 3D (Fig 1C, S1A-C): colocalising (<64 nm), adjacent (65-120 nm), and separated (>120 nm). The colocalising category included cases where the two proteins perfectly overlap in 3D, as well as cases where they partially overlap in 3D. Quantification of each configuration indicated that the two proteins most commonly colocalise in both the trachea (43,34%, n=60 vertices from 6 embryos) and the epidermis (42,23%, n=135 vertices from10 embryos). Among these colocalising cases, most of them corresponded to a complete overlap (75% of cases in the trachea and 61,54% of cases in the epidermis). We also observed a high percentage of tAJs where the two proteins are merely adjacent (33,3%, in trachea, n= 60 vertices from 6 embryos, and 34,07% in epidermis n=135 vertices from10 embryos), or even completely separated (23,3% in trachea, n= 60 vertices from 6 embryos, and 23,7% in epidermis n=135 vertices from10 embryos) (Fig 1D).

**Figure 1.**
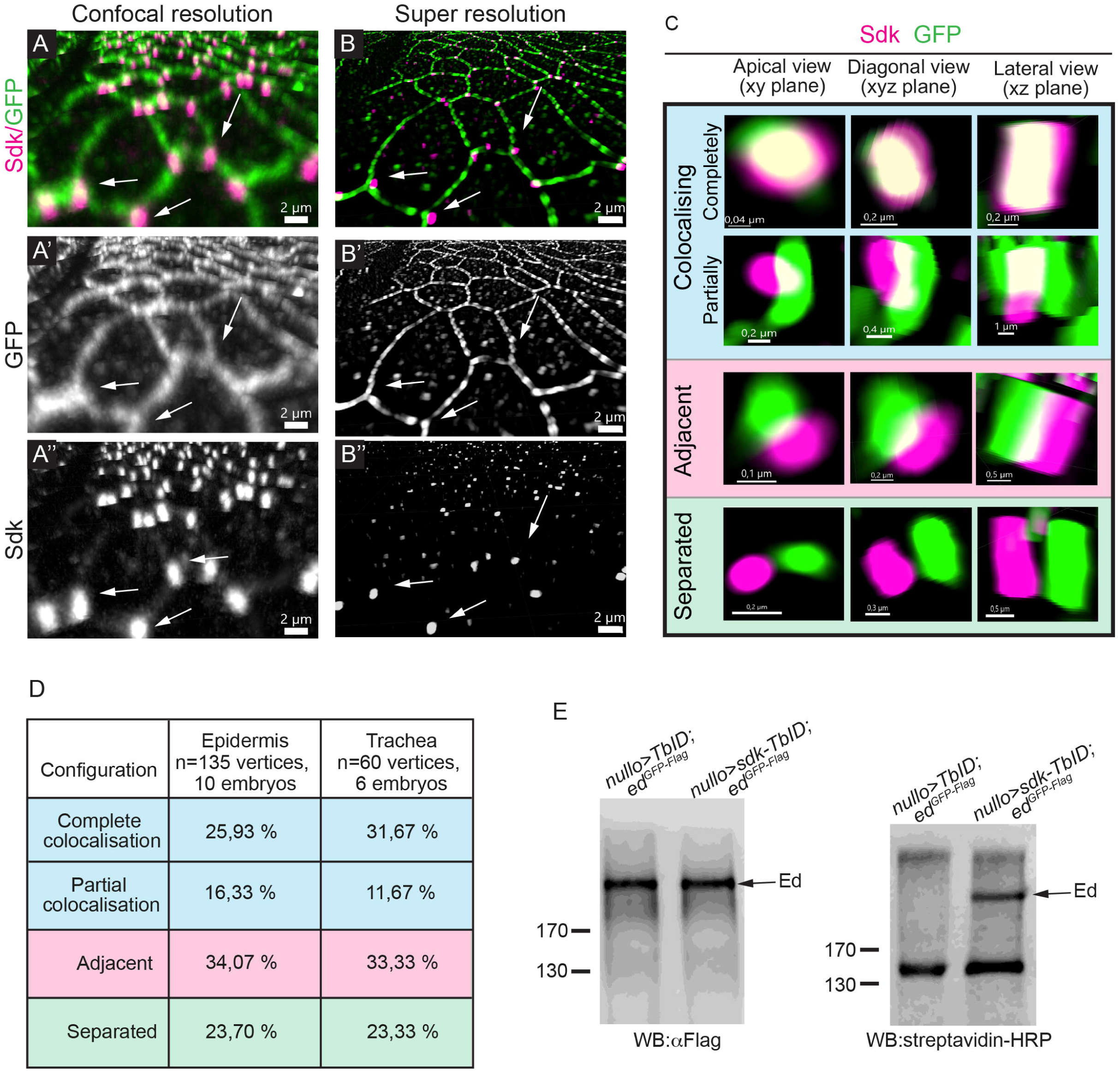
Sdk and Ed localisation at AJs. (A,B) Images of the embryonic epidermis of ed^GFP^ embryos at stage 14 corresponding to projections at conventional confocal resolution (A) and super-resolution using the SRRF-mode (B). Embryos were stained with GFP to reveal Ed accumulation (green, grey) and with an antibody against Sdk to visualise Sdk accumulation (magenta, grey). White arrows indicate points where Sdk and Ed colocalises at confocal resolution, but appear only closely positioned, rather than overlapping, under super-resolution. (C) Shows examples of the variety of configurations between Ed and Sdk in 3D. (D) Quantification of Ed/Sdk configurations in the epidermis or tracheal vertices. (E) Protein extracts from embryos carrying the *ed^GFP-Flag^*allele and expressing either *TurboID* or *sdk-TurboID* were subjected to immunoprecipitation with α-Flag antibody and analyzed by western blot using α-Flag to detect immunoprecipitated Ed protein (left panel) and streptavidin-HRP to detect biotinylated Ed protein (right panel). MW markers (in kDa) are indicated. Scale bars are indicated.

The high frequency of Sdk and Ed colocalisation suggests that these proteins may be able to directly interact or be part of the same protein complex. However, numerous instances where the two proteins appear clearly separated challenge the hypothesis that the two proteins form a stable complex. The findings would fit in a model in which Ed and Sdk engage in transient and dynamic interactions, which could be either direct or mediated by common partners. For example, direct interactions of Sdk and Ed with Cno (10, 19) could position them within a shared protein complex, given that Cno is capable of oligomerisation (25, 26). In this scenario, Sdk and Ed would remain in close proximity within a common complex linked by Cno (and/or other shared partners), and dissociation of either IgCAM protein from Cno would separate them.

### 2. Sdk and Ed in a shared protein complex

Because our results suggested possible transient interactions, we decided to perform proximity labelling assays based on TurboID that facilitates the capture of protein associations that are unstable, weak, or highly dynamic (27). We generated a UAS-Sdk-HA transgene tagged in its intracellular domain with the biotin ligase TurboID. We found that this fusion protein localised similarly to the UAS-Sdk-HA, which we had previously shown to be a functional protein capable of rescuing the *sdk* mutant phenotype (12) (Fig S2A,B). We expressed *UAS-sdk-TurboID-HA* in a general pattern using the *nulloGal4* line in embryos carrying the *ed^GFP^*allele (which is also tagged with Flag). After biotin administration, we purified the biotinylated proteins by binding to streptavidin-coated beads and probed for the presence of Ed using an anti-Flag antibody. We detected a very faint band at the expected molecular weight for the tagged Ed protein (around 200 kDa, corresponding to 160 kDa of Ed (Islam *et al*., 2003) plus the tags) (Fig S2C). To increase the reliability of this result, we optimised the experimental setup. Under the same experimental conditions, we first immunoprecipitated Ed and then tested whether it was biotinylated. We detected a band that was reproducible and reliable (Fig 1E). Altogether, these results suggest that Sdk and Ed are in close proximity, supporting the hypothesis of a direct interaction or their presence in a shared protein complex.

### 3. Sdk modulates Ed levels and its enrichment at tAJs

As we observed that Sdk and Ed largely colocalise at AJs and can potentially interact, we asked whether they regulate each other’s accumulation.

We first tested whether *sdk* modulates Ed by using the *ed^GFP^*allele and analysing the trachea and the epidermis of embryos at stage 14. Both a visual inspection (Fig 2A-F) and a subsequent quantitative analysis of Ed fluorescence intensity at both tAJs and bAJs (Fig 2G,H) showed that *sdk* affects the overall levels of Ed accumulation. We found an increase in Ed levels in the absence of *sdk*, and a decrease under *sdk* overexpression conditions, in both the trachea and epidermis when compared with control. Remarkably, the same modulation was observed at both the bicellular and tricellular junctions, despite the fact that Sdk is predominantly enriched at the tAJs (12).

**Figure 2.**
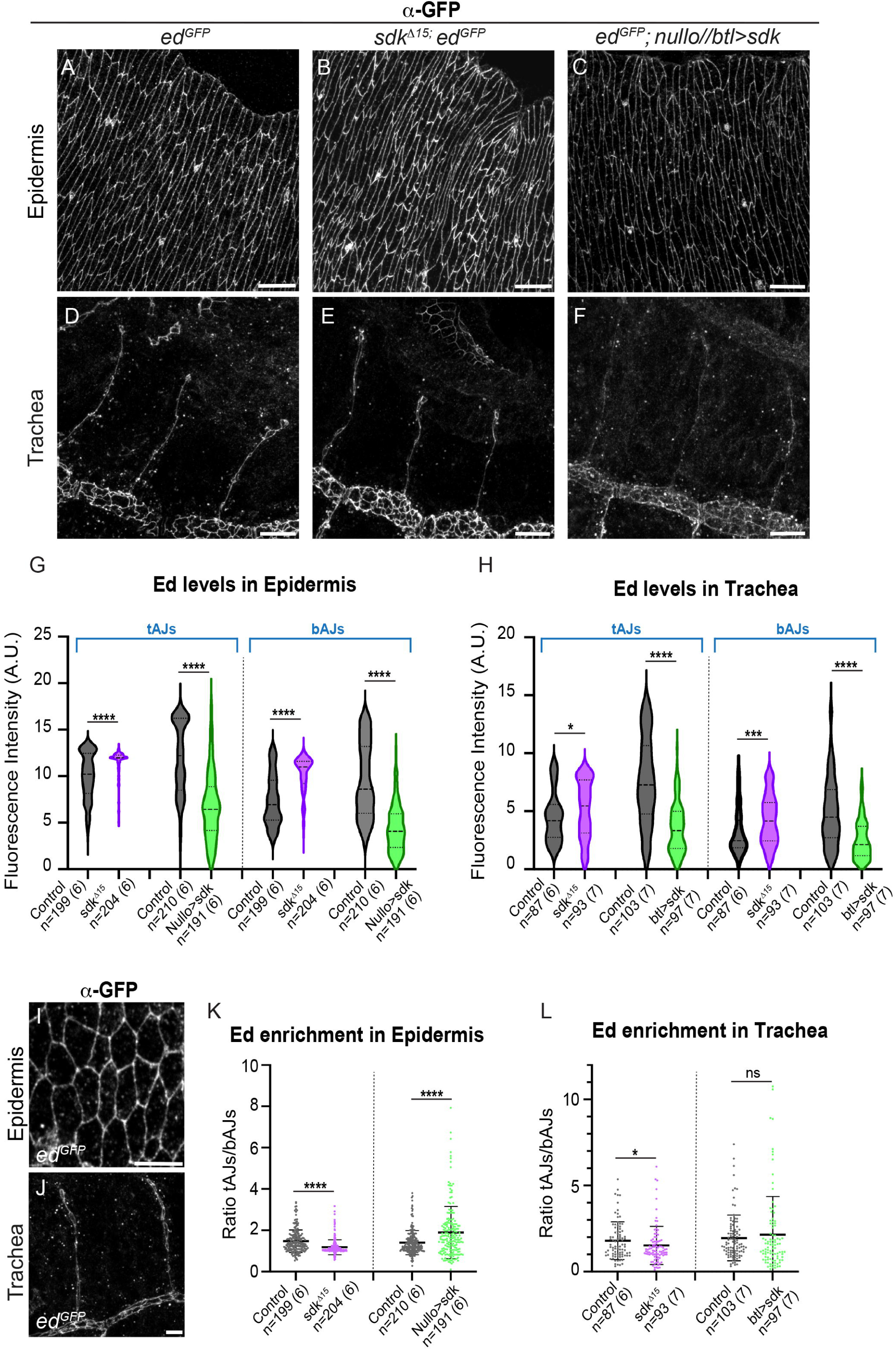
Ed accumulation in *sdk* mutant conditions. (A-F,I,J) Confocal projections of the epidermis and trachea of embryos of the indicated genotypes at stage 14 stained for GFP to reveal Ed accumulation. (G,H) Violin plots showing Ed fluorescence intensity levels both at tricellular and bicellular AJs in the different genotypes indicated, in trachea and epidermis. (K,L) Scatter plots showing the enrichment of Ed at tAJs in the epidermis and trachea and in different *sdk* mutant conditions. n=number of junctions analysed in the epidermis or tracheal dorsal branches in n=number of embryos (in brackets). Black asterisks refer to the significance with respect to control. p-values obtained by a non-parametric Mann-Whitney two-tailed test. ****p < 0.0001, 0.0001 < ***p < 0.001, 0.001 <**p < 0.01, *p < 0.05, ns not significant Scale bars A-F 10 μm; I,J 5 μm

We then analysed whether *sdk* affects the distribution of Ed protein along the AJs. In the control we found that while Ed consistently localised along the entire AJ, it was slightly enriched at tAJ. The enrichment, calculated as the ratio of tAJs levels/bAJs levels, showed an average of around 1,4-2 in epidermis and trachea in the controls (Fig 2I-L). This enrichment decreased in *sdk^Δ15^* null mutants and increased in *sdk* overexpression conditions (Fig 2K,L), indicating that *sdk* modulates the modest enrichment of Ed at tAJs.

Our results indicate an inverse correlation between the enrichment of Ed at tAJs and its overall levels both in bAJs and tAJs upon *sdk* manipulations. This is explained by a stronger effect of *sdk* on Ed levels at bAJs. Although an effect at bAJs may seem unexpected given the predominant localisation of Sdk at tAJs, several explanations could account for this observation. On the one hand, Sdk could be present at bAJs at levels that are difficult to be detected due to its strong enrichment at tAJs. This is consistent with evidence that Sdk is able to localise at bAJs in different contexts (12, 13), indicating that its localisation is not strictly exclusive to tAJs. In this scenario, Sdk could compete with Ed for plasma membrane binding or occupancy within the protein complex. Thus, the loss of Sdk would lead to increased Ed levels, due to a competition-based mechanism. Alternatively, Ed levels in the different AJs domains could be interdependent. In this case, Sdk, localised and acting mainly at tAJs, could affect Ed levels along the entire AJ through various mechanisms, including modulation of intracellular trafficking and recycling, changes in lateral membrane diffusion, or stabilisation via protein complexes. These hypotheses are not mutually exclusive.

### 4. Sdk and Ed localisation in vesicles

In the course of our analysis of the subcellular localisation of Sdk and Ed, we noticed the presence of Ed in intracellular vesicles (Fig 3A). This pattern was also observed using an antibody against the protein (Fig 3B), indicating that it reflects the normal Ed accumulation. We found that a proportion of Ed positive vesicles were also positive for Sdk. This was observed both in the epidermis (Fig 3C,E) and the tracheal system (Fig 3D,E), but the proportion of Ed vesicles that were positive for Sdk was significantly higher in the trachea (81%) than in the epidermis (23%). This suggests tissue-specific crosstalk between Sdk and Ed. In this context, it is relevant to mention that Sdk is actively involved in cell intercalation, a process that takes place in the trachea at stage 14 and that involves important remodelling of AJs (see below).

**Figure 3.**
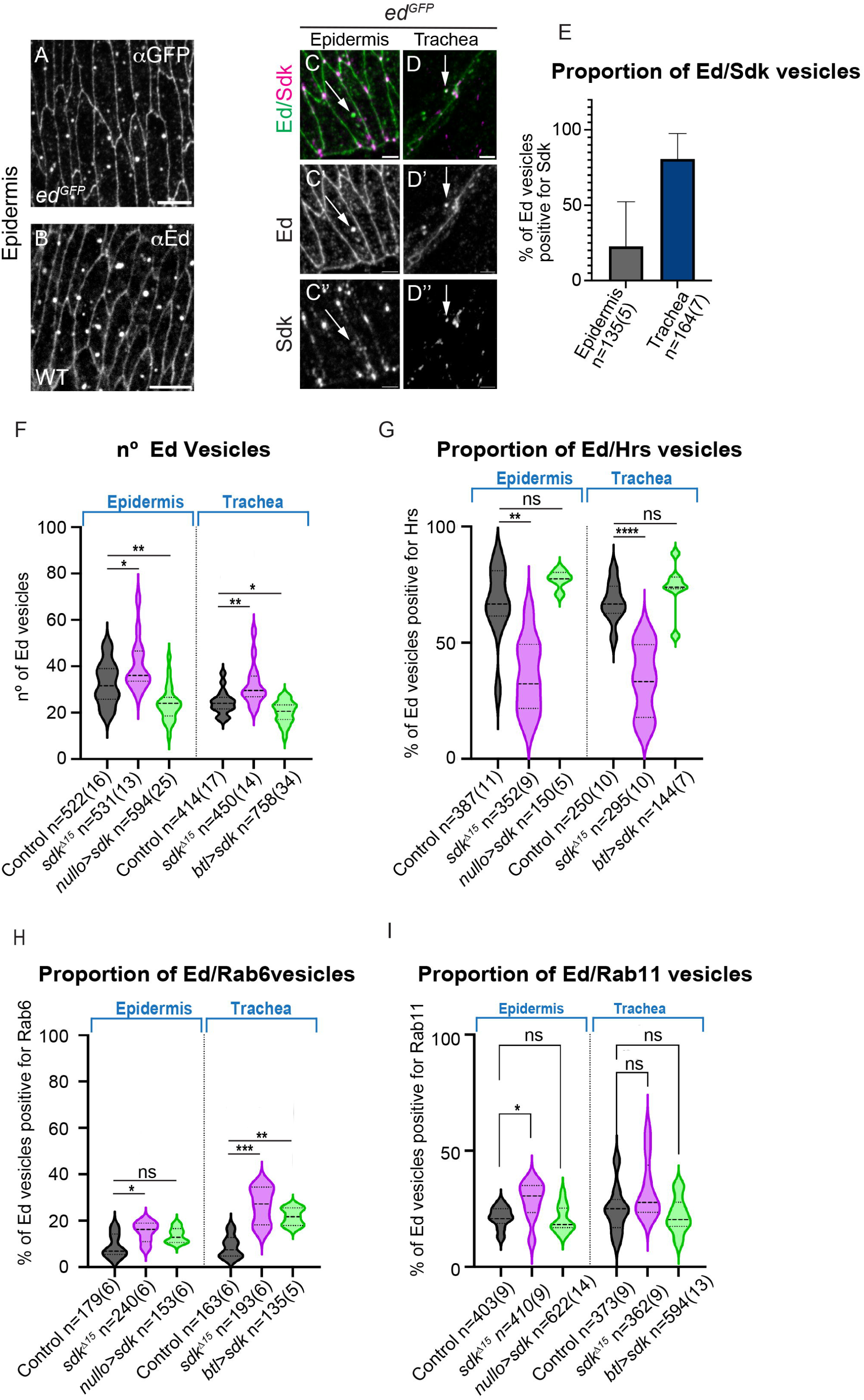
Ed intracellular trafficking. (A-D) Confocal projections of the indicated genotypes and tissues. Ed localises at AJs and also in intracellular vesicles. A proportion of Ed vesicles are also positive for Sdk in the trachea and epidermis (arrowheads in C,D). (E) Quantification of the proportion of Ed vesicles positive for Sdk in trachea and epidermis. (F-I) Violin plots with the proportion of Ed vesicles positive for the indicated markers, obtained from embryos at stage 13-14 in the indicated genotypes. Asterisks refer to the significance with respect to control. p-values obtained by ordinary one-way ANOVA with Dunnets’s multiple comparisons test. ****p < 0.0001, 0.0001 < ***p < 0.001, 0.001 <**p < 0.01, *p < 0.05, ns not significant n=number of vesicles analysed in the epidermis or tracheal dorsal branches in n=number of embryos (in brackets) Scale bars A,B 5 μm; C,D 2 μm.

### 5. Sdk affects Ed intracellular trafficking

The observation that Ed and Sdk colocalise in intracellular vesicles and that *sdk* modulates Ed levels, prompted us to investigate whether Sdk affects the pattern of vesicular Ed. Actually, we detected an increase in the total number of Ed vesicles in *sdk* mutant conditions and a decrease in *sdk* overexpression conditions, both in the epidermis and the trachea (Fig 3F). The change in the number of Ed vesicles by *sdk* manipulations correlates with the modulation of Ed levels: increased Ed levels are associated with a higher number of Ed vesicles in the absence of *sdk*, whereas reduced Ed levels correlate with fewer Ed vesicles in *sdk* overexpression conditions. This correlation between Ed vesicle number of Ed levels suggests that *sdk* may regulate Ed levels, at least in part, by modulating its intracellular trafficking.

We used various markers to determine the nature of Ed-containing vesicles under wild type, *sdk* loss of function and *sdk* overexpression conditions. On the one hand, we found that a significant part of the Ed vesicles in the wild type, both in the epidermis and trachea, were endocytic, as more than >50% of them were also positive for the ESCRT-0 complex component Hrs, a marker of late endosomes (28). This confirms the endocytic nature of a significant part of Ed vesicles (29). Interestingly, the proportion of Ed vesicles also positive for Hrs significantly decreased in *sdk* mutants and mildly increased in *sdk* overexpression conditions (although this increase did not appear to be statistically significant) (Fig 3G, S3). On the other hand, in *sdk* mutants we detected an increased proportion of Ed exocytic vesicles as revealed by the co-staining with Rab6 (Fig 3H, S3), which localises within the trans-Golgi network (TGN) and regulates trafficking from the Golgi to other membrane targets (30–32). Moreover, in *sdk* mutants we detected a mild increase in the recycling of Ed back to the membrane (significant only in the epidermis), as shown by co-staining with Rab11, a marker of recycling from the recycling endosome to the membrane (33). Conversely, a mild decrease in recycling was observed under *sdk* overexpression conditions, although it did not reach statistical significance (Fig. 3I, S3).

Altogether our results indicate that *sdk* modulates the intracellular trafficking of Ed and are consistent with a model in which it does so by promoting endocytosis while hindering trafficking to the membrane. In this model, the decrease in endocytosis together with the increase in recycling and exocytic trafficking observed in *sdk* mutants would account for increased Ed levels. We note, however, that the results under *sdk* overexpression conditions do not always perfectly align with this model and are not always complementary to those observed for loss of function. However, overexpression experiments should be interpreted with caution, because overexpressed proteins may produce unintended artefactual effects. We suggest that *sdk* may act indirectly in traffic regulation, possibly by modulating other partners that control trafficking, such as actin regulators (34, 35), and/or by directly interacting with and modulating endocytic/exocytic factors on its way to the membrane and during recycling, and/or even by competing directly with Ed for occupancy within trafficking vesicles. Such mechanisms could explain a role for *sdk* in modulating Ed accumulation not only at tAJs but also at bAJs, despite not being localised at bAJs.

### 6. Sdk accumulation in gain and loss of *ed* conditions

After analysing the impact of *sdk* in Ed accumulation, we then analysed the impact of *ed* in Sdk accumulation. To completely deplete both the maternal and zygotic activity of *ed (ed^1X5^ m+z)* we generated *ed^1X5^* germ line clones (*ed^1X5^*, an amorphic allele, (36)). While these mutant embryos displayed strong morphogenetic defects in various organs, including the trachea and epidermis (37, 38), Sdk was still able to localise at the tAJs in both tissues (Fig 4A,B,E,F). However, the severe morphogenetic defects did not allow a reliable evaluation of Sdk levels, preventing a proper assessment of whether *ed* affects Sdk accumulation.

**Figure 4.**
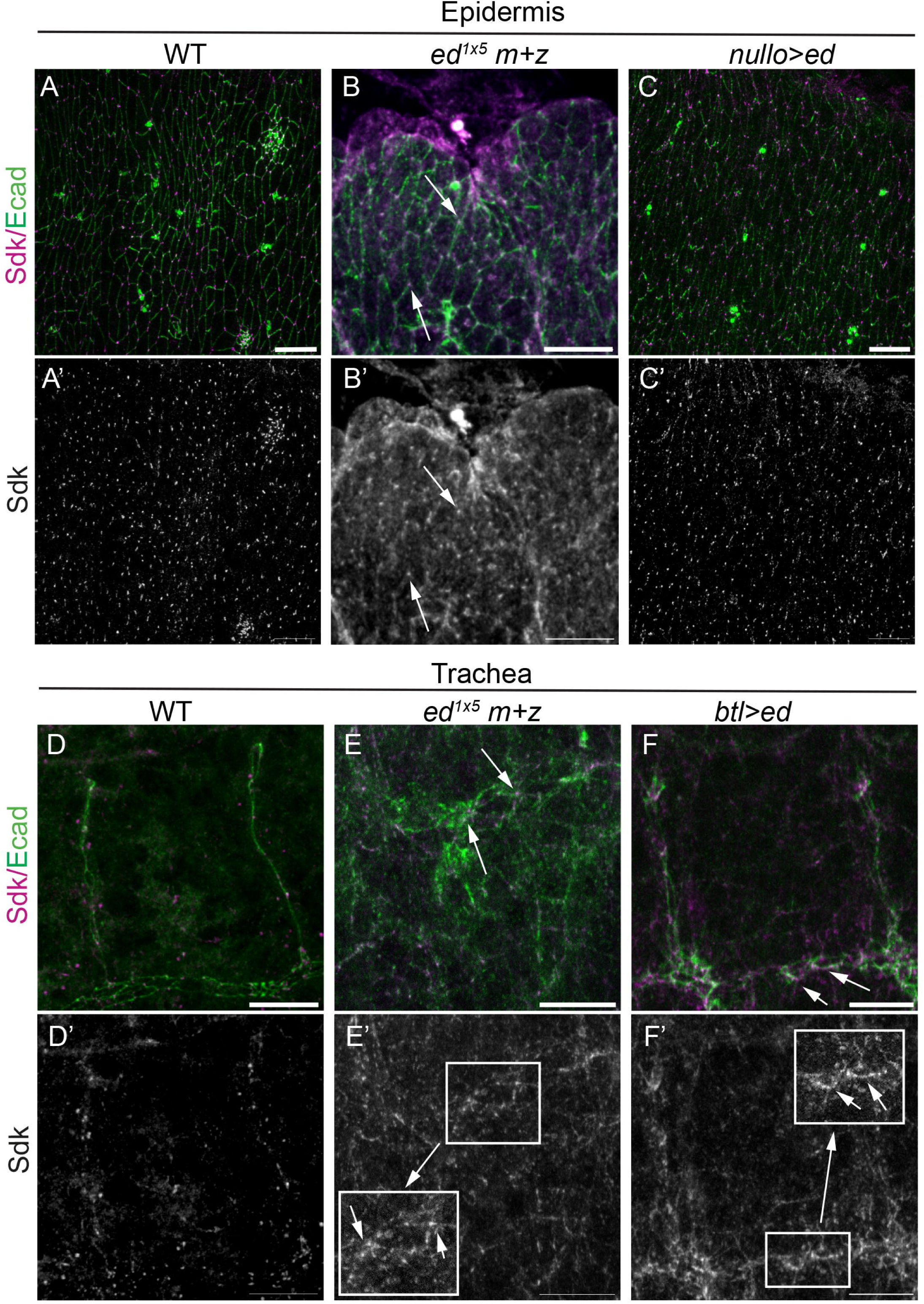
Sdk localisation in *ed* manipulation conditions. Confocal projections of the indicated genotypes and tissues of embryos at stage 14 stained for Sdk (magenta, grey) and Ecad (green). (A-C) In the epidermis, Sdk localises at tAJs in *ed* maternal+zygotic mutants (arrows in B). In *ed* overexpression conditions Sdk is enriched at tAJs but also displays some expansion towards the bAJs in some cells (arrows in C). (D-F) In the trachea, the absence of maternal+zygotic *ed* activity leads to strong defects, but Sdk can still localise at tAJs (arrows in E,E’ and inset). When *ed* is overexpressed, Ecad appears more diffuse and we detect an expansion of Sdk towards the bicellular junctions (arrows in F,F’ and inset). Scale bars 10 μm.

Thus, we resorted to an overexpression assay using an available *UAS-ed* line (36). In the epidermis, Sdk remained largely enriched at tAJs when *ed* was overexpressed, as in controls. However, we noticed a mild expansion of Sdk accumulation towards the bicellular junction in some cells (Fig 4C, S4A-B). Quantification of Sdk accumulation at tAJs and bAJs indicated a clear increase in Sdk levels at bAJs, and a more moderate increase at tAJs in *ed* overexpression conditions. As a result, Sdk becomes slightly less enriched at tAJs (tAJs/bAJs ratio) when *ed* is overexpressed (Fig S4C-E). In the trachea, the overexpression of *ed* resulted in a stronger expansion of Sdk towards the bicellular junctions than in the epidermis (Fig 4F), which may point again to tissue-specific crosstalk between these two proteins that could relate to the cellular events taking place (e.g. cell intercalation in the trachea). However, we cannot rule out the possibility that these differences are due to the variable effectiveness of the Gal4 driver used.

Taken together, while our findings indicate that Sdk is still able to localise at tAJs in *ed* null mutants, the overexpression experiments suggest also an effect of *ed* in Sdk accumulation. This effect could be direct, for instance by Ed recruiting Sdk into the complex, or indirect, for instance mediated by Ecad. Actually, we detected a decrease in Ecad in *ed* overexpression conditions in the trachea (Fig S4D,E) and Ecad has been shown to regulate Sdk localisation (12). Additionally, *ed* activity is known to affect tension (16, 37) and tension is known to regulate Sdk localisation (12). Therefore, effects on Ecad and/or tension caused by *ed* overexpression may indirectly regulate Sdk localisation.

### 7. Sdk and Ed genetically interact

To explore a potential functional relevance of the Ed/Sdk cross-regulation we analysed various *sdk/ed* genetic combinations, focusing specifically on tracheal cell intercalation and on the embryonic morphology, which have been reported to be affected by *sdk* and *ed* mutants respectively.

Tracheal cell intercalation drives the rearrangement of cells in most tracheal branches and ensures the transformation of intercellular AJs into autocellular ones (39). To quantify the intercalation defects in different mutant conditions, we used our established protocol that categorises dorsal branches into four classes based on the severity of the defects (40). Class I represents fully intercalated branches, while class IV corresponds to branches completely lacking intercalation (Fig S5A). As we previously reported (12), *sdk* mutants exhibit pronounced intercalation defects (Fig 5A,B,F). In contrast, tracheal intercalation has not been previously evaluated in *ed* mutants. To analyse this, we first considered using embryos completely lacking *ed* (*ed^1X5^ m+z*), but their severe embryonic defects precluded a reliable assessment of tracheal intercalation. We therefore analysed embryos lacking only the zygotic contribution of *ed* (*ed^1X5^* z). Although this condition does not represent a full loss of *ed* activity, it allows embryonic development to proceed sufficiently for a systematic evaluation of the tracheal pattern. Interestingly, we observed a previously undescribed, very mild effect in tracheal intercalation, which was less severe than that seen in s*dk* mutants (see Fig 5C,F). Taking into account that the presence of maternal contribution under these conditions is likely masking a much greater requirement for *ed* during the intercalation process, this observation shows a role for *ed* in tracheal intercalation. We then analysed the tracheal intercalation pattern under different genetic combinations of *sdk* and *ed*. We first tested transheterozygous conditions by removing one copy of each gene. This combination resulted in a very mild phenotype that was not significantly different from the control (Fig S5B,C,E). We then analysed double mutants for *sdk* and e*d*, which showed an extreme effect on intercalation, stronger than in any mutant condition alone (Fig 5D,F). The severe effect suggests a synergistic epistasis between *sdk* and *ed*, possibly revealing cooperative functions. The overexpression of *sdk* did not produce tracheal intercalation defects (Fig S5D). We then tested whether *sdk* overexpression could rescue the mild intercalation defects produced by the absence of zygotic *ed* and we found that this was the case (Fig 5E,G), suggesting that *sdk* can substitute at least in part for *ed* activity during tracheal intercalation. Altogether, our results reveal that both *ed* and *sdk* participate in the process of cell intercalation.

**Figure 5.**
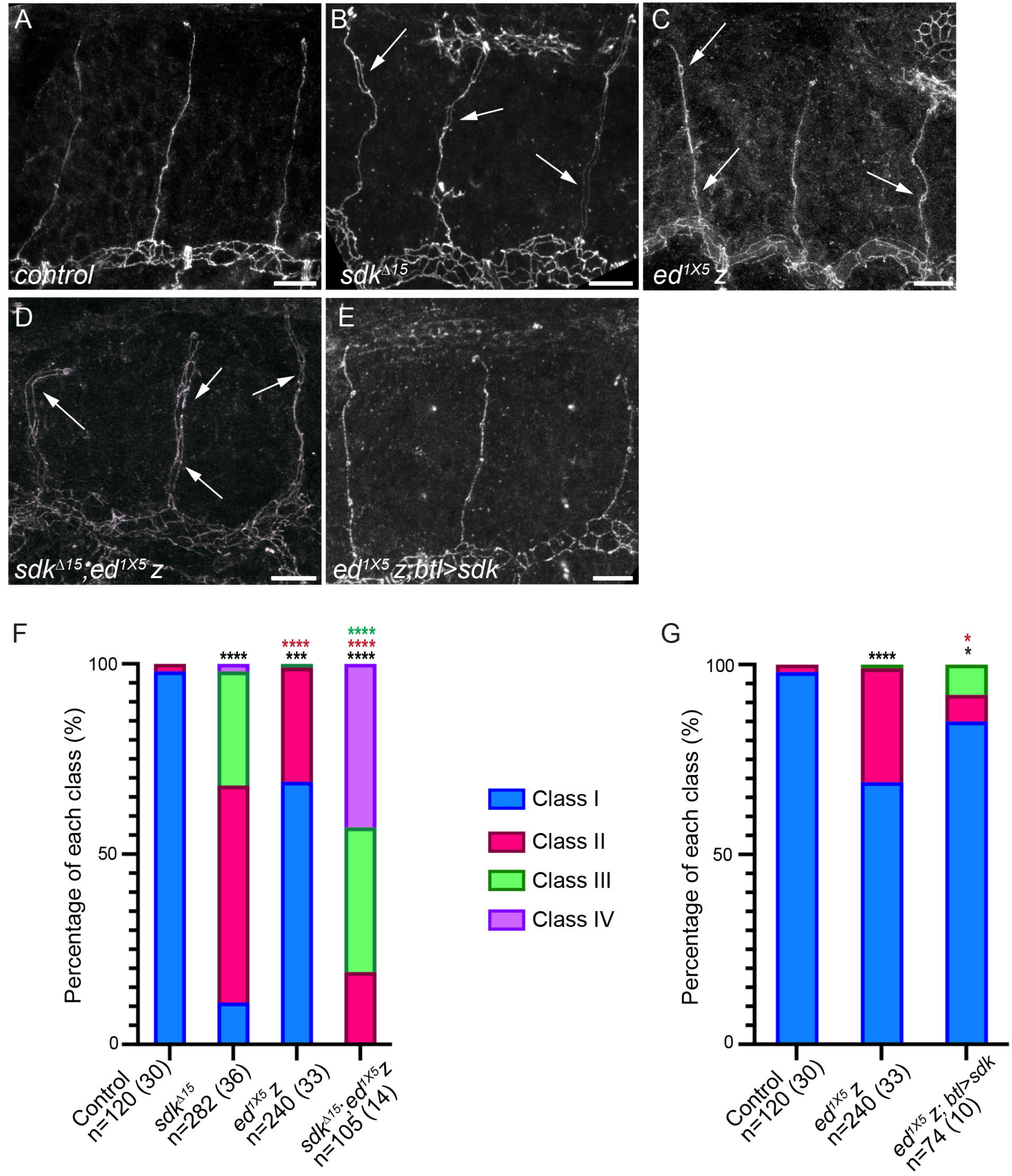
Analysis of tracheal intercalation. (A-E) Confocal projections of stage 15/16 embryos stained for Ecad to mark AJs in the tracheal branches in the indicated genotypes. White arrows point to intercalation defects. (F,G) Quantification of the percentage of branches of each class of intercalation defect for each genotype. . n = number of branches analysed from n=number embryos (in brackets) analysed. p-values were obtained using the Kruskal–Wallis test with Dunn’s multiple comparisons test. ****p < 0.0001, 0.0001 < ***p < 0.001, 0.001 <**p < 0.01, *p < 0.05. Black asterisks indicate comparison with first column (control), red asterisks with second column (*sdk* in F and *ed* in G), and green asterisks with third column (*ed* in F). Scale bars 10 μm.

We then analysed the overall embryonic morphology in different *sdk/ed* combinations. To quantify the defects in a systematic manner, we focused on dorsal closure and head development, as these events have been demonstrated to be regulated by *ed* (16, 17, 37). In wild type conditions, stage 16/17 embryos have completed dorsal closure exhibiting a straight dorsal midline, and the head region is internalised (Fig 6A,S6A,B) after undergoing a process of head involution (41). We classified the defects into 4 classes (Fig 6): 1) the “very mild” class, characterised by the presence of minor defects observed in the dorsal midline, which appears wiggly or not completely straight (Fig 6B,S6C,D), 2) the “mild” class, characterised by incomplete dorsal closure (Fig 6C,S6E,F), 3) the “intermediate” class, exhibiting defects both in dorsal closure and head deformation (Fig 6D,S6G,H), and the “severe” class, typified by the presence of strong head deformations, strong dorsal closure defects, and generalised deformation and/or holes in the embryo (Fig 6E,S6I,J). Our analysis revealed that *sdk* mutants exhibited previously unreported defects in dorsal closure, which we classified as very mild (Fig 6F). In contrast, zygotic *ed* mutants displayed a spectrum of phenotypes from mild to intermediate, primarily characterised by defects in dorsal closure (Fig 6F). Double mutants displayed a markedly severe phenotype (Fig 6F), indicating that loss of *sdk* exacerbates the zygotic *ed* phenotype. As observed in the case of tracheal intercalation, the severe defects suggest a synergistic epistasis between *sdk* and *ed.* Finally, the overexpression of *sdk* did not rescue the zygotic *ed* mutant phenotype, as we observed in the trachea, and instead it resulted in a severe phenotype (Fig 6G), suggesting that *sdk* cannot substitute for *ed* activity during embryonic patterning.

**Figure 6.**
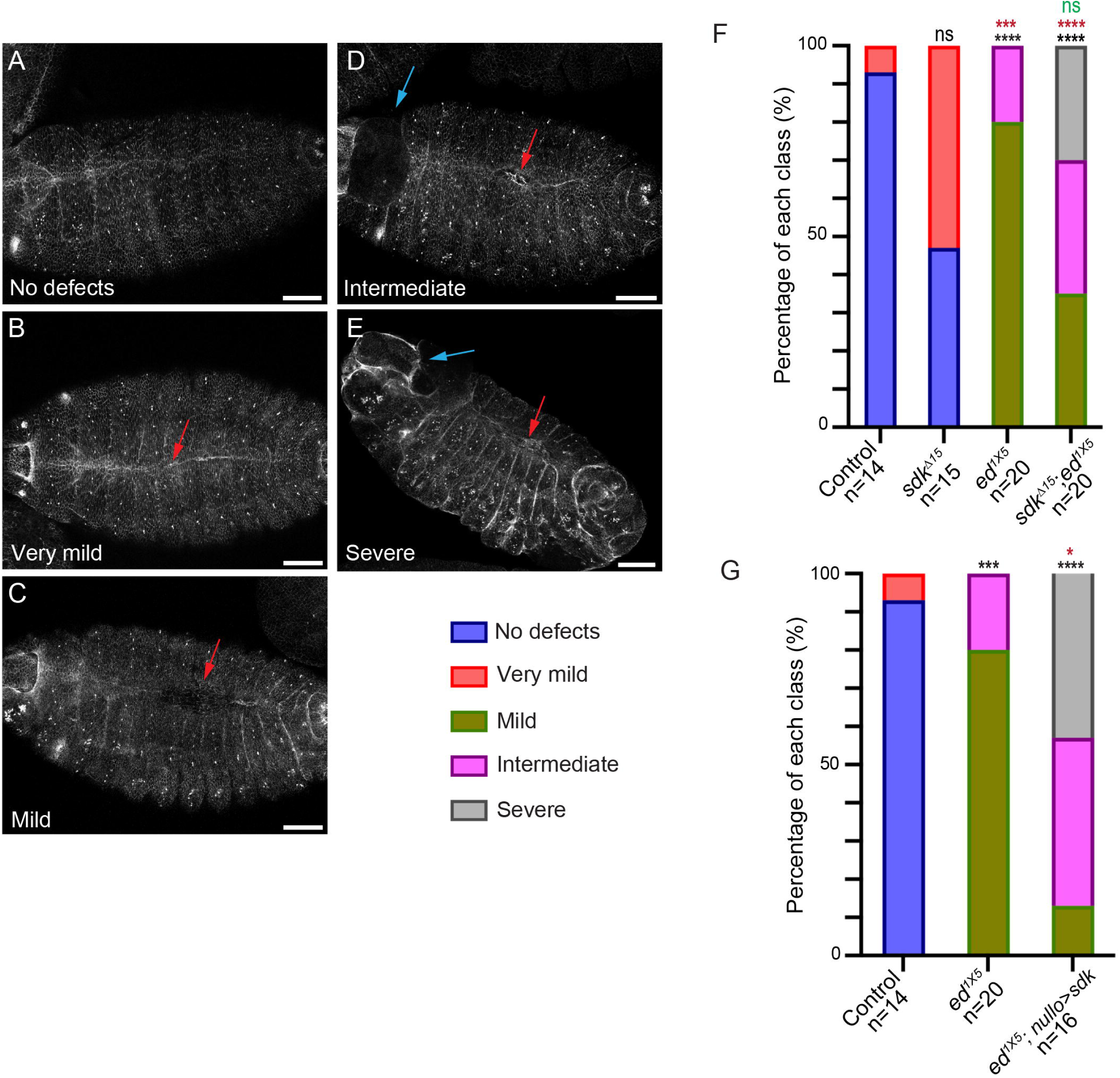
Analysis of embryonic morphology. (A-E) Confocal projections of dorsal views of stage 15/16 embryos stained for Ecad to mark AJs. Examples of the different categories of defects in dorsal closure (red arrows) and head (blue arrows) are shown. (F,G) Quantification of the percentage of embryos with each class of defects for each indicated genotype. n = number of embryos analysed. p-values were obtained using the Kruskal–Wallis test with Dunn’s multiple comparisons test . ****p < 0.0001, 0.0001 < ***p < 0.001, 0.001 <**p < 0.01, *p < 0.05, ns not significant. Black asterisks indicates comparison with first column (control), red asterisks with second column (*sdk* in F and *ed* in G), and green asterisks with third column (*ed* in F) Scale bars 30 μm.

Our findings reveal several points. First, they indicate that both *sdk* and *ed* contribute to dorsal closure and tracheal cell intercalation, which had not been previously recognised. Second, our results indicate that each morphogenetic event relies more heavily on one of the two factors. This may reflect the specific cellular behaviours occurring in each process, with dorsal closure requiring cell elongation and tracheal intercalation requiring cell rearrangement. In this context, *sdk* appears more directly involved in cell repositioning, likely by regulating tension at tAJs. In contrast, *ed* may be better suited to regulate the formation of the supracellular actin cable required for dorsal closure and for cell elongation, consistent with its localisation along the entire AJ (16, 17). Finally, the genetic interactions observed point to a functionally significant relationship between the two genes, suggesting cooperative activities. This could be consistent with a model in which both proteins localise to, and interact within, a shared complex, contributing to its function by modulating the activity of downstream effectors in a context-specific manner.

### 8. Sdk and Ed converge and regulate Pyd and Cno

We previously showed that Sdk binds to Pyd and Cno, and that Sdk promotes the enrichment of these proteins at tAJs (10, 12). We suggested that Pyd and Cno would link Sdk to actin cytoskeleton to maintain cellular tension and allow vertex remodelling during tissue rearrangements. As it is the case for Sdk, Ed has been shown to also bind Cno (19). Additionally, Cno and Pyd have been shown to work together to support cell junction stability (42).

Because both *sdk* and *ed* genetically interact and may regulate common intracellular adaptor proteins that control epithelial organisation, we investigated whether they act together to regulate the accumulation of Cno and Pyd. We measured the tAJs enrichment (ratio tAJs/bAJs) of these proteins in the epidermis at different developmental stages under different *sdk/ed* combinations.

As we previously showed (12), our results indicated that Sdk regulates Cno tAJs enrichment. In *sdk* mutants, both at stage 10 or later at stage 13/14, the enrichment decreased compared to the control (Fig 7A,C,E, S7A,C,E). The lack of zygotic *ed*, instead, had no impact on Cno enrichment at tAJs (Fig 7A,B,E, S7A,B,E). As expected, *sdk/ed* double mutants also clearly affected the tAJs enrichment in a similar way as in *sdk* mutants (Fig 7A,D,E, S7A,D,E).

**Figure 7.**
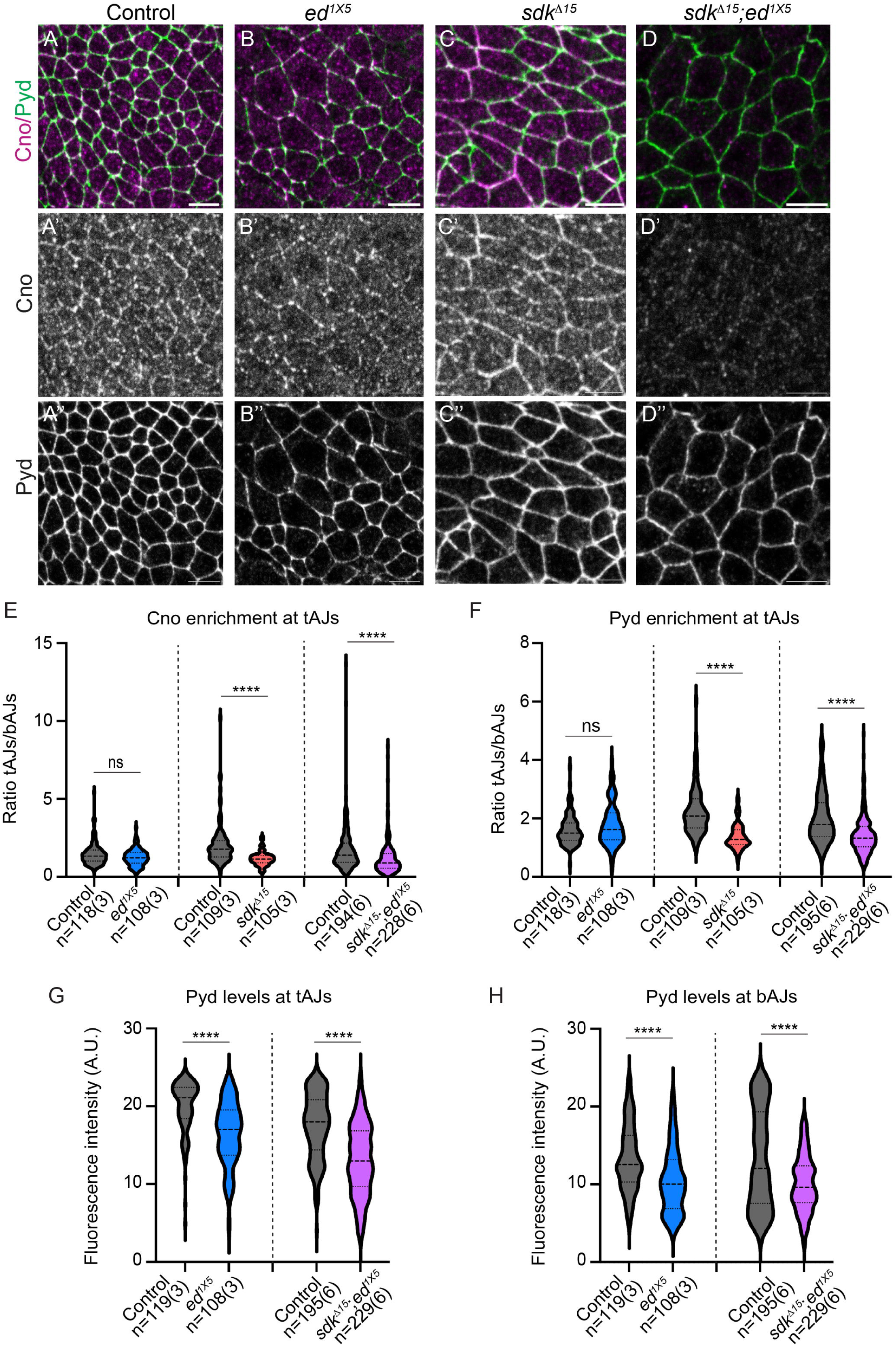
Analysis of Cno and Pyd accumulation. (A-D) Confocal projections of the epidermis of stage 10 embryos stained for Cno (magenta, grey) and Pyd (green, grey) in the indicated genotypes. (E,F) Violin plots with the quantification of the enrichment at tAJs of Cno and Pyd in the different conditions. (G,H) Violin plots with the quantification of Pyd fluorescence intensity levels both at tricellular and bicellular AJs in the genotypes indicated. n = number of junctions analysed from n=number embryos (in brackets) analysed. p-values were obtained using the non-parametric Mann–Whitney two-tailed test. ****p < 0.0001, 0.0001 < ***p < 0.001, 0.001 <**p < 0.01, *p < 0.05, ns not significant. Scale bars 5 μm.

In the case of Pyd, we found that, as previously described (12), the tAJs enrichment clearly decreases in the absence of *sdk* at st 10 (Fig 7A,C,F). We note, however, that at later stages, this Pyd tAJs enrichment does not seem to strongly depend on *sdk* (FigS7A,C,F). This suggests that different mechanisms may operate at different time points to enrich Pyd at tAJs, such as redundant pathways or stabilisation through interactions with proteins other than Sdk. Similar to the case of Cno, we observed that the lack of zygotic *ed* activity does not have an impact on Pyd tAJs enrichment (Fig 7A,B,F, S7A,B,F). The absence of both *sdk* and *ed* leads to a clear decrease of tAJs enrichment at early stages (Fig 7A,D,F) but not at later stages (Fig S7A,D,F), in line with the findings with *sdk* mutants alone.

We noticed, however, that while *ed* did not significantly affect the tAJs enrichment of Pyd, it clearly affected the general levels of this protein (Fig 7A,B, S7A,B). We quantified the intensity of Pyd levels and we detected a clear and significant decrease both at tAJs and bAJs (Fig 7G,H). This effect on the general levels of Pyd was also observed in *sdk ed* double mutants (Fig 7A,D,G,H, S7A,D). We analysed whether *ed* also affected the general levels of Cno. We could not detect an effect at early stages, but we observed a decrease on Cno levels at tAJs and bAJs at later stages (Fig 7B, S7A,B,G). An effect of *ed* on Cno levels could be expected considering that Cno was shown to bind Ed (19). The absence of an effect at early stages could reflect a specific temporal regulation of Cno by *ed*, or the presence of sufficient maternal *ed* to maintain Cno levels. Alternatively, this temporal specificity may reflect the complex interplay and molecular interactions among the different components of a shared protein complex.

Altogether, our results indicate that *sdk* and *ed* modulate the levels and distribution of both Cno and Pyd. *sdk* primarily modulates the enrichment of these adaptor proteins at the tAJs, consistent with its enrichment there. Additionally, we find a previously undescribed regulation of Pyd by *ed* along the entire AJ. The interactions of Sdk and Ed with Pyd and Cno that we identify here, together with those previously described, suggest that Pyd and Cno contribute significantly to the functional requirements of Ed and Sdk during key developmental processes, including dorsal closure, head morphogenesis, and cell intercalation (42–44).

Based on the results here reported and on previous observations (10, 12, 19, 42), we propose a model in which large protein complexes form both at bAJs and tAJs, incorporating a mixture of both common (e.g. Ed, Cno, Pyd) and junction-specific/enriched components (e.g. Sdk). Within these complexes, numerous and distinct molecular interactions occur simultaneously between different components, and their dynamics are likely influenced by the specific composition of each complex, for example, whether the complex belongs to a tAJ or a bAJ (and thus have more or less Sdk). These interactions can affect the composition and behaviour of bAJs and tAJs complexes in a cross-regulatory manner, through mechanisms such as lateral diffusion of components within the membrane, intracellular trafficking and recycling, transient associations that contribute to signaling and structural regulation, or protein stabilisation. Moreover, individual proteins within the complex may recruit additional partners, thereby conferring further specificity, robustness, and functional versatility to the junctional architecture. Together, these layers of interactions likely give rise to a highly intricate regulatory environment that enables epithelial junctions to remodel efficiently during morphogenetic events, maintaining both the robustness necessary for tissue integrity and the flexibility required for precise cell rearrangements such as intercalation.

## ACKNOWLEDGEMENTS

We thank A. Letizia for contributions to the initial stages of this project. We thank the IBMB Molecular Imaging Platform for technical help. We acknowledge the Bloomington Stock Centre and the Developmental Studies Hybridoma Bank for fly lines and antibodies.We thank J. Treisman, J. Modolell, A. Carmena, L. Nilson and T. Nakamura for kindly providing fly lines and antibodies. We thank the members of the Llimargas and Casanova labs and S. Araújo for helpful discussions, and A. Letizia, J. Casanova and J. Treisman for critical reading of the manuscript.

This work was funded by the Spanish Ministerio de Ciencia e Innovación to Marta Llimargas (MICINN, https://www.ciencia.gob.es/va/), grants numbers PGC2018-098449-B-I00, PID2021-126689NB-I00 and PID2024-157683NB-I00). The funders had no role in study design, data collection and analysis, decision to publish, or preparation of the manuscript. The authors declare no competing interests.

## Author contributions

Conceptualization: ML

Methodology: FC, MLE, ML

Validation: FC, MLE, ML

Investigation: JC, MLE

Writing – original draft: ML

Writing –review & editing: MLE, ML

Supervision: ML

Funding acquisition: ML

## Data and resource availability

All relevant data and resources can be found within the article and its supplementary information. Data reported in this paper and resources will be shared upon request by the lead contact ML.

## Declaration of generative AI and AI-assisted technologies in the writing process

During the preparation of this work, Microsoft Copilot was used exclusively to assist with language editing and to improve clarity and readability. Following the use of this tool, the authors critically reviewed and revised the content as necessary and assume full responsibility for the final version of the published article.

## MATERIALS AND METHODS

### Drosophila strains

All *Drosophila* strains (Table 1) were maintained at 25°C under standard laboratory conditions. Overexpression and RNA interference (RNAi) experiments were performed using the Gal4/UAS system (Brand and Perrimon, 1993), with crosses maintained at 29°C to maximize expression. The Gal4 drivers employed were btl-Gal4 (for the expression of the desired transgenes in all tracheal cells) and nullo-Gal4 (for the expression in the epidermal cells). The mutant chromosomes were balanced over FM7 or CyO, marked with LacZ or GFP, in order to facilitate the recognition of the homozygous mutant embryos.

**Table 1.**
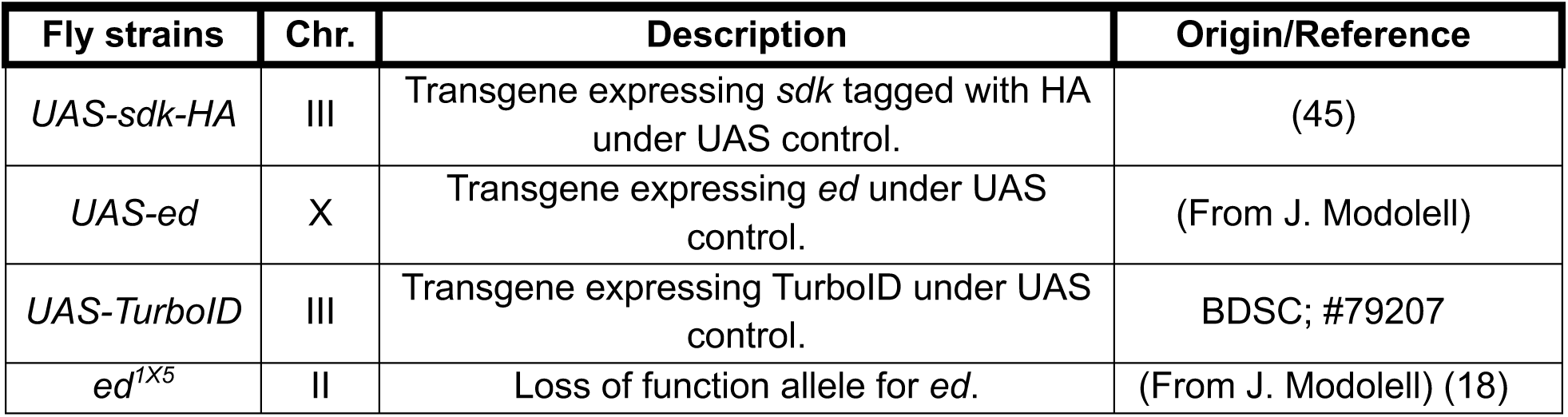
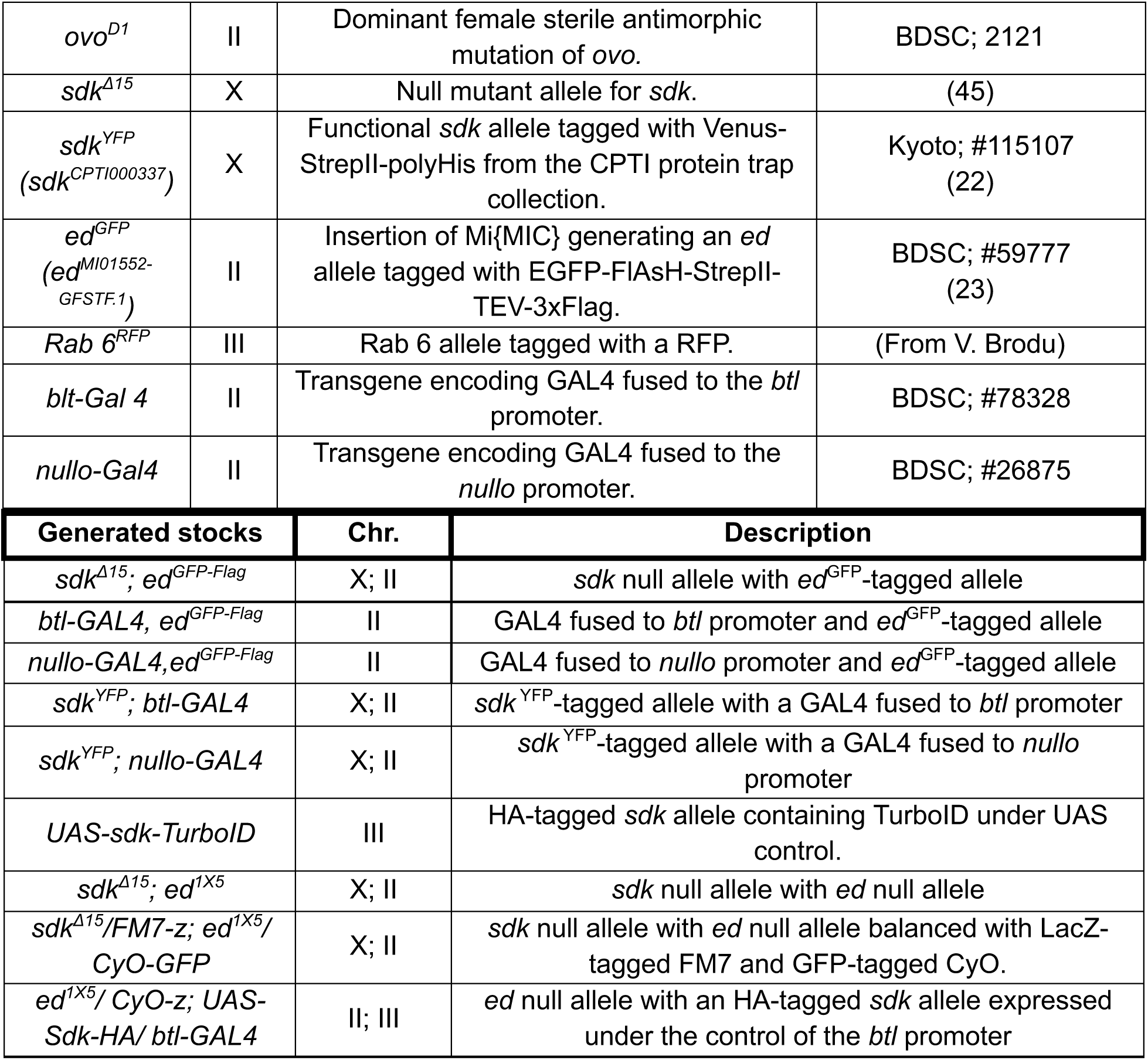
Fly strains used during the study. Abbreviations used: Chr: Chromosome number; BDSC: Bloomington *Drosophila* Stock Center. VDRC: Vienna *Drosophila* Resource Center.

### Generation of germ-line mutant clones using FLP/FRT *ovo^D1^* system

To remove *ed* maternal and zygotic contribution, we generated *ed^1X5^* germ line clones using Dominant Female Sterile (DFS) technique (46). Heterozygous females for *ed^1X5^* carrying a FRT sequence (*hs-Flp; ed^1X5^*FRT40A*/CyO lacZ*) were mated to males carrying the dominant female sterile mutation, *ovo^D1^* FRT40A/*CyO*. Second- to third-instar larval progeny was subjected to two 45-minutes heat shocks at 37 °C in a water bath, with a two-hour recovery period at room temperature. Following eclosion, adult females heterozygous for *ed^1X5^* and *ovo^D1^*were crossed with *ed^1X5^* FRT40A/*CyO lacZ* males. Only embryos that were homozygous for the mutation, resulting from the recombination, continued developing and were collected for immunostaining.

### Embryo fixation and immunostaining

Embryos collected from agar plates for a period of 24 hours were dechorionated with 100% commercial bleach for 2 minutes and rinsed with Triton 0.1%. The embryos were collected and fixed for 20 minutes (10 minutes for ECad staining) in 4% formaldehyde, PBS (0.1 M NaCl, 10 mM phosphate buffer, pH 7.4) and heptane 1:1 to generate holes in the vitelline membrane of the embryos. After removing the bottom phase of the fixation solution, the embryos were washed three times with methanol and used for immunostaining. The fixed embryos were rinsed 3 times, for 20 minutes each wash, with PBT (PBS with 0.3% Triton X100) with 0.5% BSA (Bovine Serum Albumin from Roche). Primary antibody (Table 2) incubation was performed in fresh PBT-BSA at 4°C overnight. The following day, the embryos were rinsed three times for 20 minutes, using PBT-BSA. Secondary antibody (Table 2) incubations were performed in the dark for a minimum of two hours at room temperature in PBT-BSA. The embryos were washed for 1 hour at room temperature in 3 washes of PBT and mounted in Fluoromount-G (Southern Biotech), with the exception of those used for super-resolution analysis, which were mounted in ProLong Glass Antifade Mountant (Thermo Fisher) or RapiClear 1.49 (SunJin Lab).

**Table 2.**
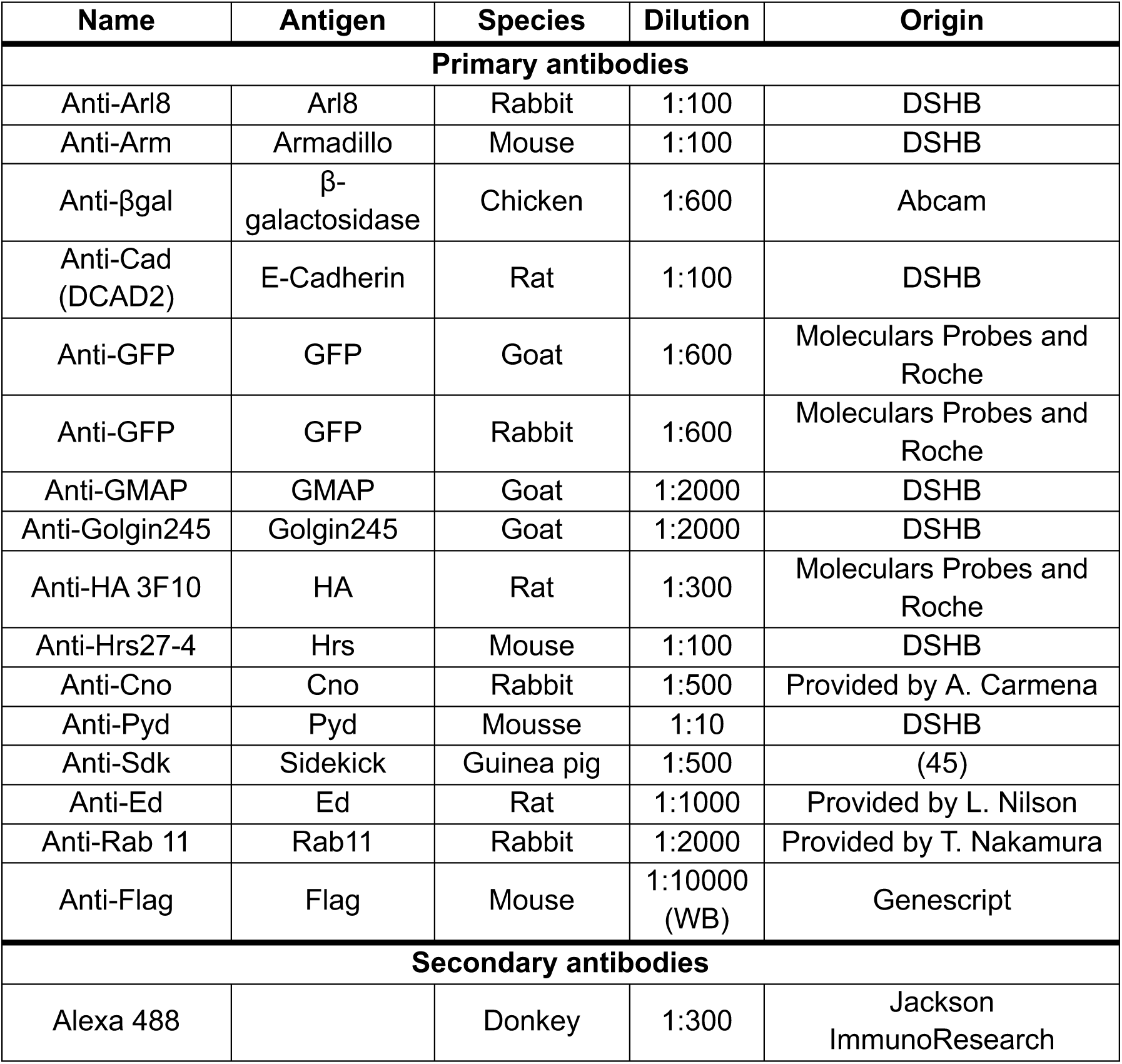
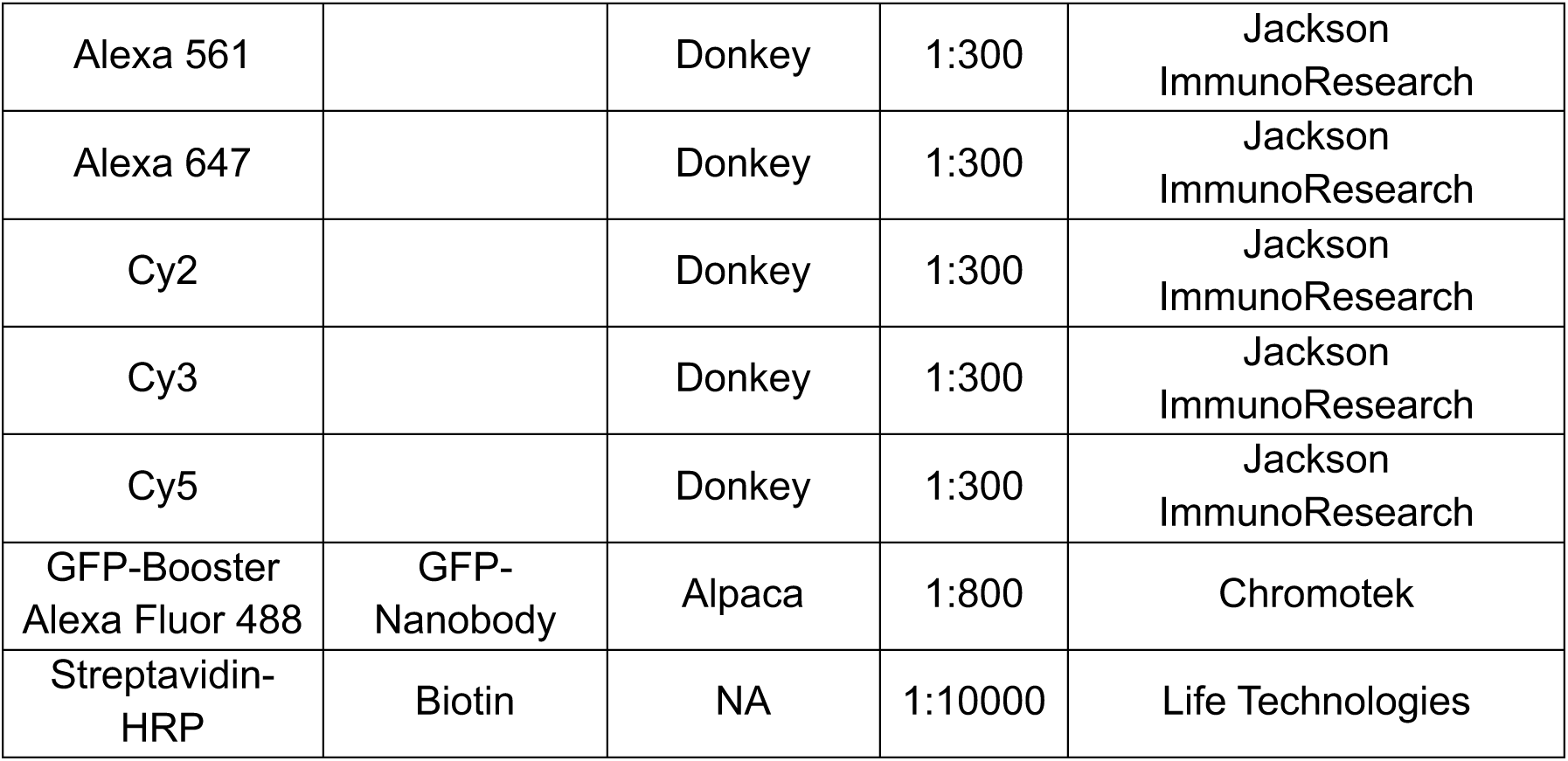
Antibodies and stains used in the study. Abbreviations used: Developmental Studies Hybridoma Bank (DSHB).

### Microscopy

Fluorescence confocal images of fixed embryos were obtained with Leica TCS-SPE system using 63x (1.40-0.60 oil) objectives. For super-resolution images a Drangofly 505 spinning disk confocal (Andor) equipped with an 100x (1.40-0.60 oil) objective and the SRRF (Super Resolution Radial Fluctuations) software-based approach for super-resolution (Gustafsson *et al*., 2016) was used.

Fiji (ImageJ) (47) was used for measurement and adjustment of the images. Unless otherwise indicated in the text, confocal images are maximum-intensity projections of Z-stack sections. Figures were assembled with Adobe Illustrator.

### Image Analysis

#### Super-resolution image analysis

Super-resolution analysis of Sdk and Ed accumulation was conducted using 2-5 z-sections from embryos at stage 13 or 14 acquired on the Dragonfly 505 microscope, using the SRRF (super-resolution radial fluctuation)-stream mode with around 700 interactions. The relative subcellular localisation of Sdk/Ed at tAJs was analysed using the line selection tool from Fiji: a line was drawn from the centre of a fluorescent point of one protein to the centre of the fluorescent point of the other protein, and the distance between the two centres was measured. Based on the different spatial configurations observed between Ed and Sdk, their relative localisation was classified into three categories—colocalising, nearby, and separated—each defined by a specific distance range (<64 nm, 65–120 nm, and >120 nm, respectively)

#### Quantification and colocalisation of vesicles

To quantify the number of vesicles and perform a colocalisation analysis, images of stage 13-14 embryos were taken and analysed manually using the Fiji circle selection tool to quantify Ed vesicles in each specific region. In the epidermis, this region included 10 individual cells which are not in direct contact with the amnioserosa. In the tracheal system, the region included two dorsal branches at the central part of the embryo, excluding the cells in direct contact with the dorsal trunk. The selected vesicles were added in the ROI Manager. For the colocalisation analysis, the colocalising vesicles of Ed vesicles with the different intracellular trafficking markers were identified manually.

#### Intensity levels

We estimated Ed, Arm, Cno, Pyd and Sdk protein levels under different *sdk* and *ed* conditions by analysing their intensity levels after immunostaining. For each experiment, we collected, fixed and stained control and experimental embryos together. The acquisition settings for intensity measurements were kept constant between control and experimental embryos within each preparation. We obtained maximum projections of confocal sections to include either 2 dorsal branches or a region of 10 epidermal cells at stage 10 or 13-14, as indicated. Integrated density levels were measured in the trachea and epidermis using the circle selection tool of Fiji, with an area of 0.04-0.09 μm², depending on the protein, for all measurements. Three distinct ROIs were analysed: 1) at tricellular junctions, 2) at the adjacent bicellular junction, and 3) at the centre of a cell contributing to the tricellular junction (as background measurement). The mean background of each embryo was subtracted from each raw intensity measurement. Enrichment at the tAJs/bAJs was calculated dividing the fluorescence intensity at each tricellular junction by the intensity at the adjacent bicellular junction. We compared the fluorescence levels of the different proteins in the trachea/epidermis and bicellular/tricellular junctions across different experimental conditions and internal control embryos using statistical analysis.

### Quantification and statistics

Data from quantifications was imported and treated in the Excel software and in GraphPad Prism 10.1.1 (GraphPad Software), where graphics were finally generated. Graphics presented in this study are violin plots (indicating median and quartiles), scatter plots (indicating mean with standard deviation) or columns (indicating the mean and the standard deviation). For statistical analysis between two groups we applied the unpaired two-tailed Student’s t-test with Welch’s correction, and the non-parametric Mann-Whitney two-tailed test when data was not normally distributed. We used the ordinary one-way ANOVA with Dunnett’s multiple comparison test for comparisons between three or more groups, and the Kruskal-Wallis test as a non-parametric test to compare differences between independent groups for the graphs. Differences were considered significant when p<0.05. Significant differences are shown in the graphs as *p < 0.05, **p < 0.01, ***p < 0.001, ****p < 0.0001. The sample size (n) is indicated in the figures or legends.

### Generation of UAS-Sdk-TurboID

We obtained *UAS-sdk-HA-TurboID* construct from *UAS-Sdk-HA* (45) by cloning the TurboID sequence in the XbaI site of Sdk. Details of the construct are available under request. Transgenic lines with this construct were obtained via phiC31-mediated integration in the “Transgenesis Service” of the “Centro de Biología Molecular Severo Ochoa” (CBM, Madrid).

### Turbo-ID proximity labelling

Crosses of *UAS-Sdk-HA-TurboID* and *nulloGal4,ed^GFP^*flies were allowed to lay eggs overnight at 25°C. Protein extracts were prepared from embryos that were lysed with a Dounce homogeniser in RIPA buffer (50 mM Tris-HCl pH8,150 mM NaCl, 0.1% SDS, 0.5% sodium deoxycholate,1% Triton X-100, 1mM PMSF and protease inhibitors (cOmplete Tablets, Roche). Protein extracts were incubated with Streptavidin Dynabeads (Invitrogen) and analysed by Western blot using anti-Flag antibodies (Genscript), the Immobilon ECL reagent (Millipore) and Odyssey M Imaging System (LICORbio).

Immunoprecipitation and proximity labelling (PL) were combined to increase resolution. Extracts were immunoprecipitated using anti-Flag antibodies (Genscript) followed by incubation with Protein G Dynabeads (Invitrogen). Immunoprecipitates were analysed by Western blot using HRP-conjugated streptavidin (Life Technologies), the Immobilon ECL reagent (Millipore) and the Odyssey M Imaging System (LICORbio).

---

**Figure S1 (related to Fig 1). Sdk and Ed subcellular localisation** (A-C) Super-resolution images using the SRRF-mode of the embryonic epidermis *of ed^GFP^* embryos at stage 14. Embryos were stained with GFP to reveal Ed accumulation (green, grey) and with an antibody against Sdk to visualise Sdk accumulation (magenta, grey). Examples of different Sdk/Ed configurations are shown: colocalising, adjacent and separated, boxed in the yellow rectangle.

Scale bars 2 μm

**Figure S2 (related to Fig 1). Arm accumulation in *sdk* mutant conditions** (A,B) Pattern of accumulation of Sdk-TurboID-HA when expressed in the epidermis compared with Sdk-HA.

C) Biotinylated proteins from cell lysates of embryos carrying the *ed^GFP-Flag^* allele and expressing either *TurboID* or *sdk-TurboID* were isolated by streptavidin beads and analysed by western blot using α-Flag antibody to detect Ed protein. Input correspond to 10% of the pull-down material. MW markers (in kDa) are indicated on the left side.

Scale bars 20 μm.

**Figure S3 (related to Fig 3). Ed intracellular trafficking** (A,B) Confocal projections of *ed^GFP^* embryos stained for GFP (green) to visualise Ed and with the vesicular marker indicated (magenta). Magnifications correspond to the vesicles encircled in the boxed regions.

Scale bars 5 μm, magnifications 1 μm

**Figure S4 (related to Fig 4). Sdk localisation in *ed* overexpression**(A,B,F,G) Confocal projections of the epidermis of stage 14 *nulloGal4-UASed* embryos. Embryos were stained for Sdk (magenta, grey) and Ecad (green, grey). (A,B) Sdk appears more disperse long the AJ in *ed* overexpression conditions (arrows in B). (C-E) Quantification of Sdk enrichment at tAJs relative to bAJs and levels of Sdk at bAJs and tAJs in nulloGal4>*UASed* embryos compared to control (*UASed*) sibling embryos. n = number of junctions analysed from n=number embryos (in brackets). p-values obtained by a non-parametric Mann-Whitney two-tailed test. ****p < 0.0001, *p < 0.05. (F,G) Confocal projections of the trachea of stage 14 *btlGal4-UASed* embryos. Embryos were stained for Sdk (magenta) and Ecad (green, grey).

Scale bars A,B 5 μm, D,E 10 μm.

**Figure S5 (related to Fig 5). Tracheal intercalation** (A) Color-coded scheme of the different categories of intercalation defects. Class I corresponds to complete intercalation (no stretches of intercellular junctions), class II corresponds to one or two short intercellular stretches, class III corresponds to two or more large intercellular stretches and class IV corresponds to no presence of autocellular stretches (no intercalation). (B-D) Confocal projections of stage 15/16 embryos stained for Ecad to mark AJs in the tracheal branches in the indicated genotypes. (E,F) Quantification of the percentage of branches of each class of intercalation defect for each genotype. n = number of branches analysed from n=number embryos (in brackets) analysed. p-values were obtained using the Kruskal–Wallis test with Dunn’s multiple comparisons test in E and the non-parametric Mann–Whitney two-tailed test in F. ****p < 0.0001, ns not significant. Black asterisks indicate comparison with first column (control).

Scale bars 10 μm.

**Figure S6 (related to Fig 6). Embryonic morphology defects** (A-E) Confocal projections of stage 15/16 embryos stained for Ecad to mark AJs. Images show details of the dorsal midline to highlight the defects in dorsal closure, and the head region of embryos shown in Figure 6

Scale bars 20 μm.

**Figure S7 (related to Fig 7). Cno and Pyd accumulation** (A-D) Confocal projections of the epidermis of stage 13 embryos stained for Cno (magenta, grey) and Pyd (green, grey) in the indicated genotypes. (E,F) Violin plots with the quantification of the enrichment at tAJs of Cno and Pyd in the different conditions. (G) Violin plots with the quantification of Cno fluorescence intensity levels both at tricellular and bicellular AJs at different embryonic stages in control and *ed* mutants. n = number of junctions analysed from n=number embryos (in brackets) analysed. p-values were obtained using the non-parametric Mann–Whitney two-tailed test. ****p < 0.0001, 0.0001 < ***p < 0.001, 0.001 <**p < 0.01, *p < 0.05, ns not significant.

Scale bars 5 μm.

