## Supplementary figures and images for "Crosstalk between Echinoid and Sidekick, two IgCAM proteins, modulates Adherens Junction dynamics and tissue remodelling"

### Supplemental Figures

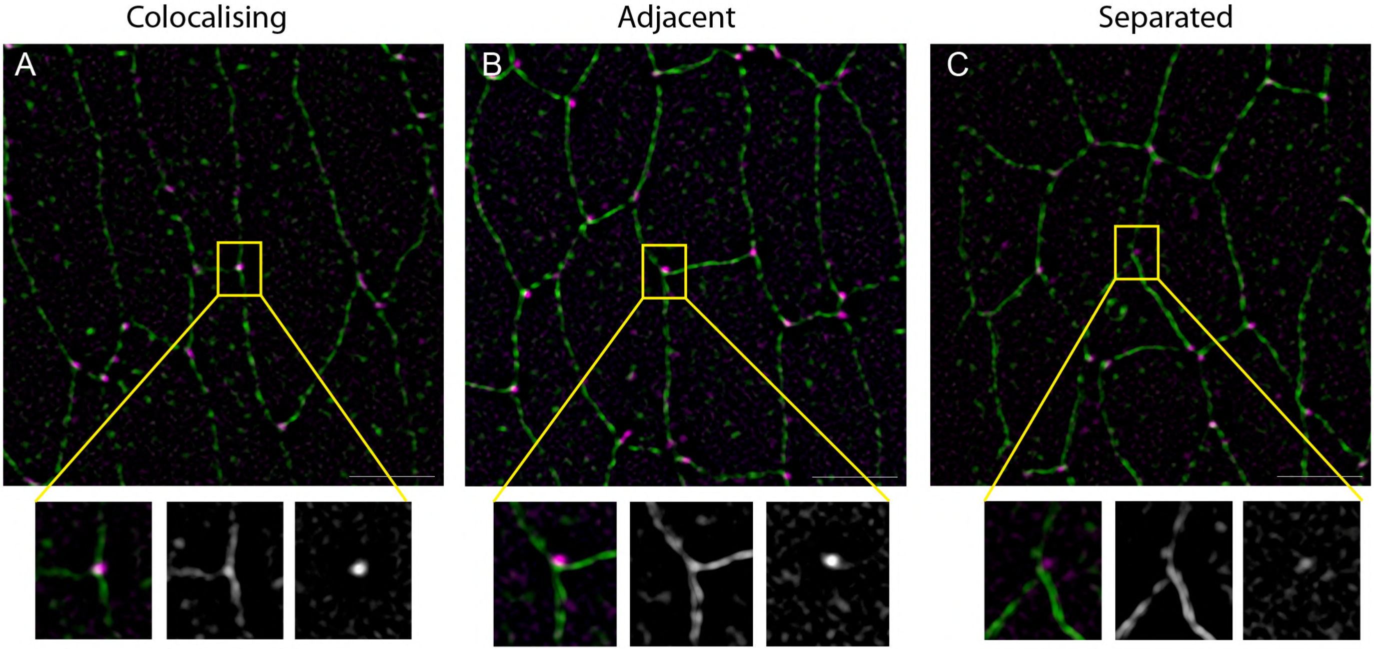


**Figure S1**

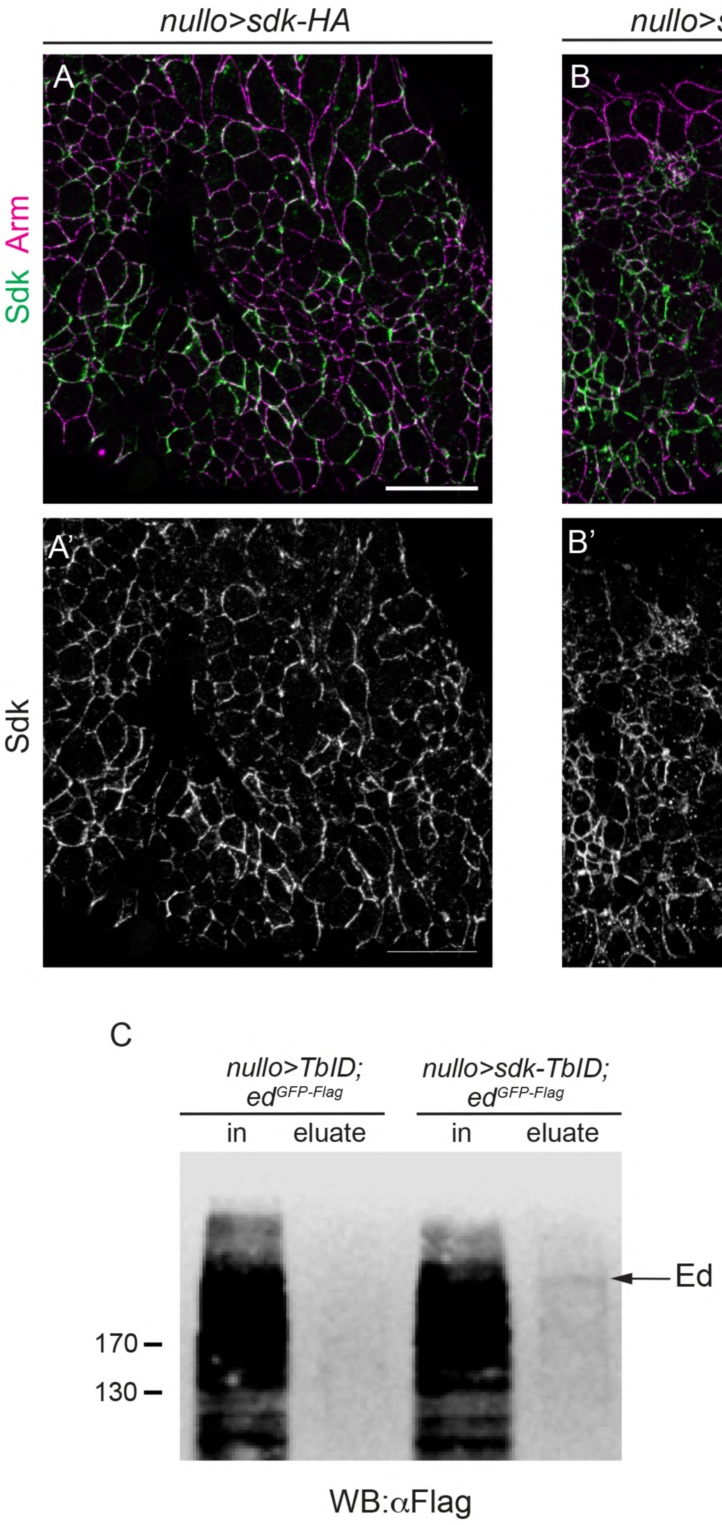

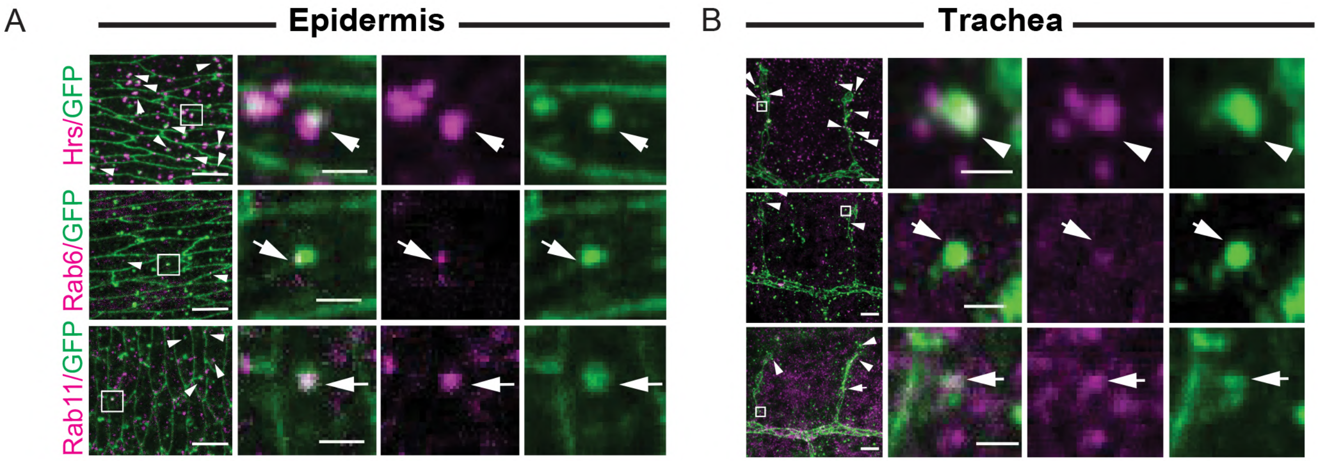


**Figure S3**

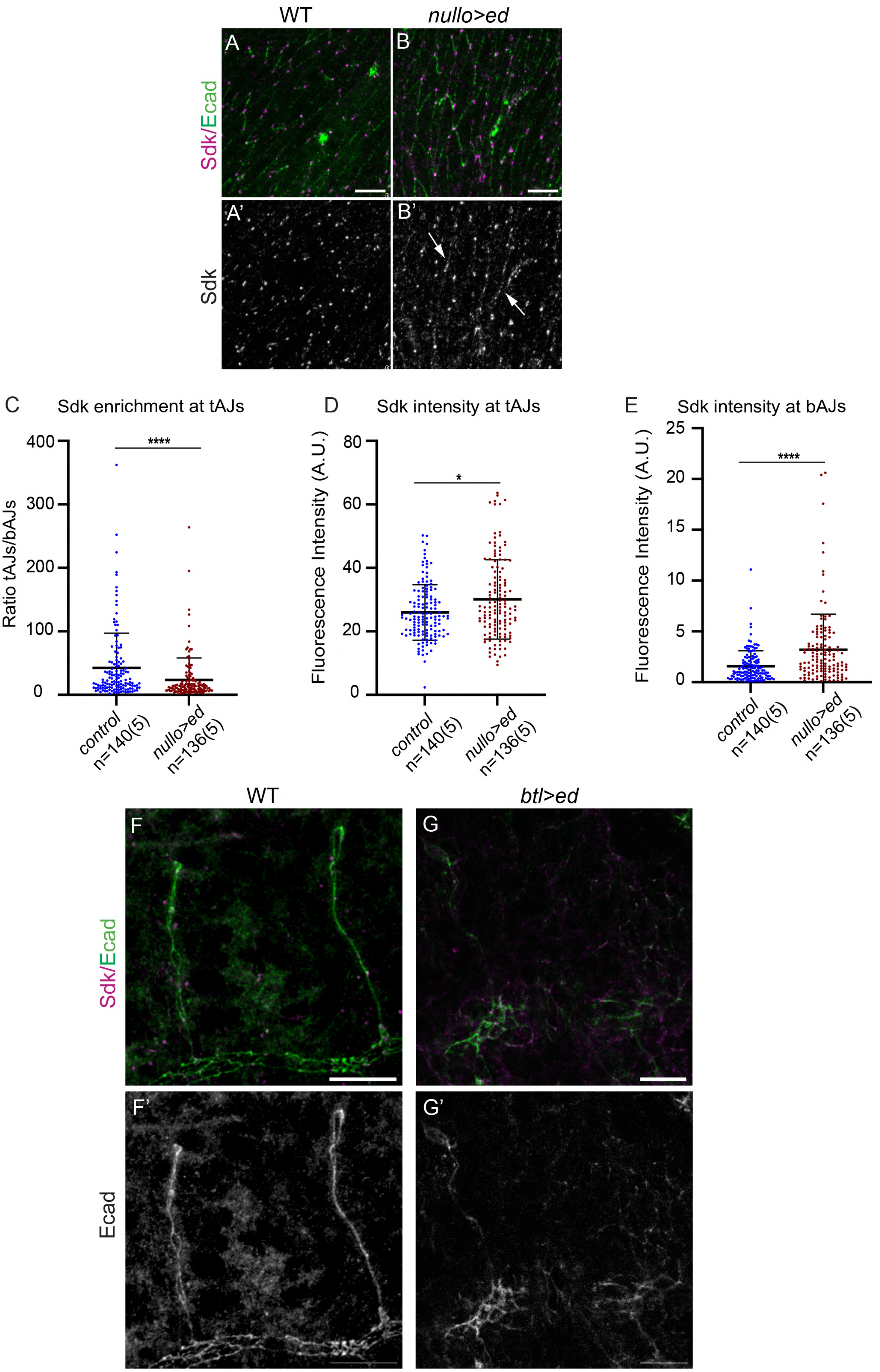

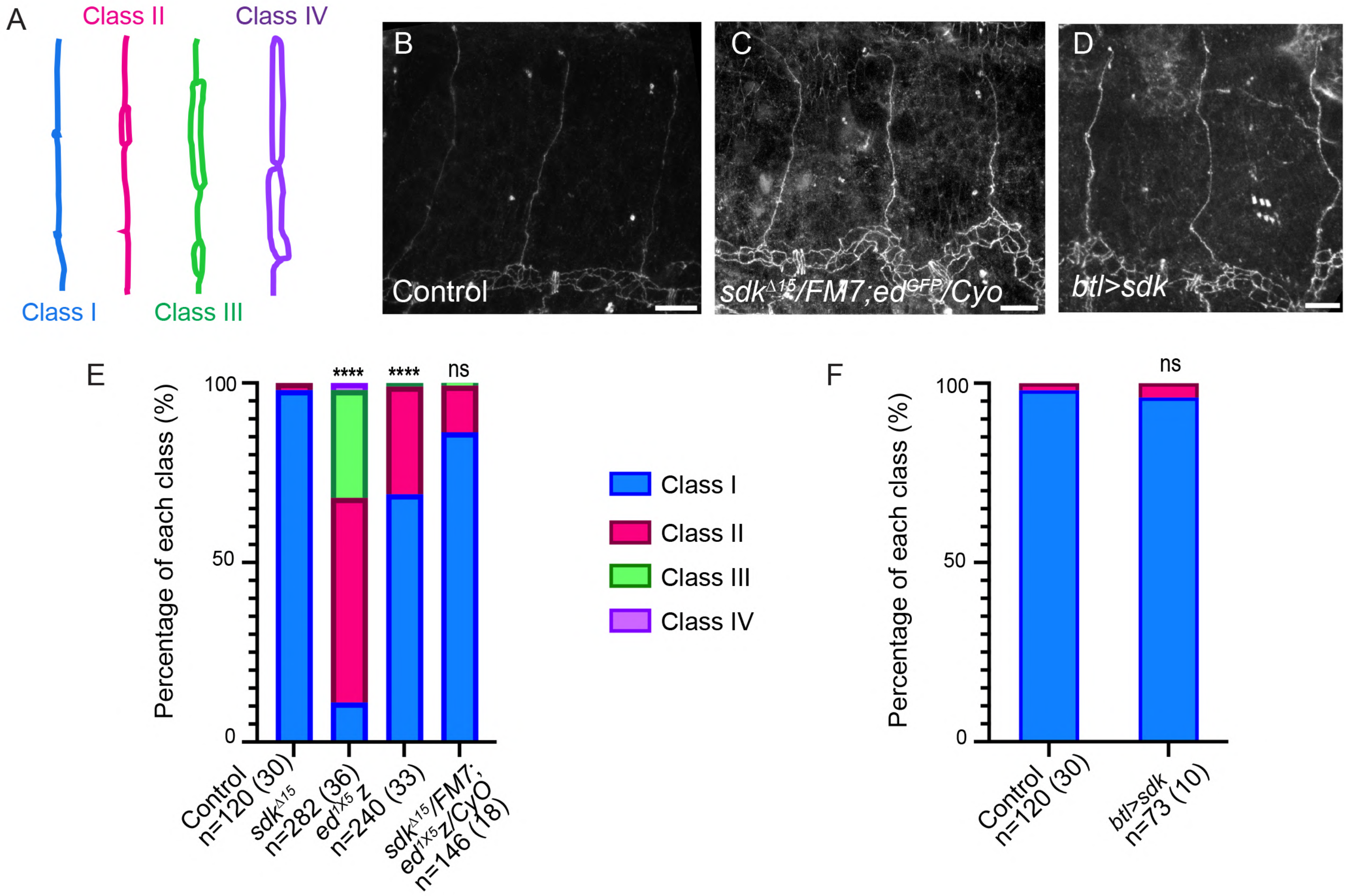

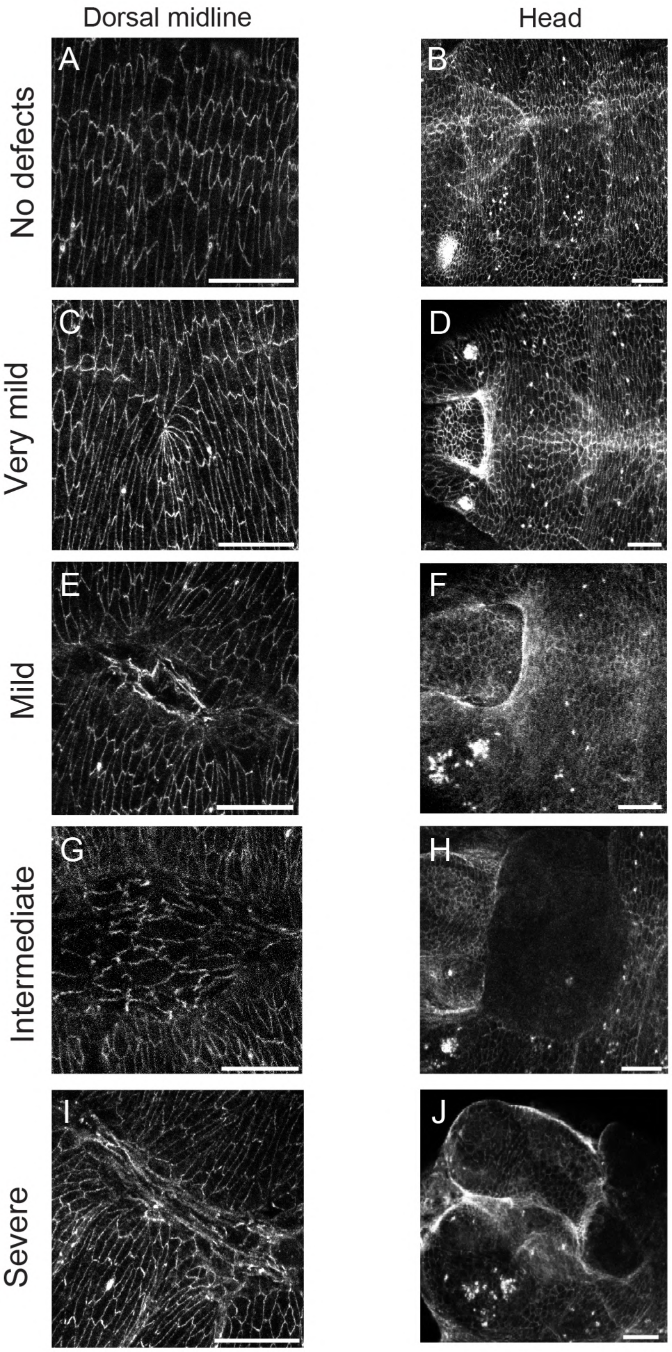

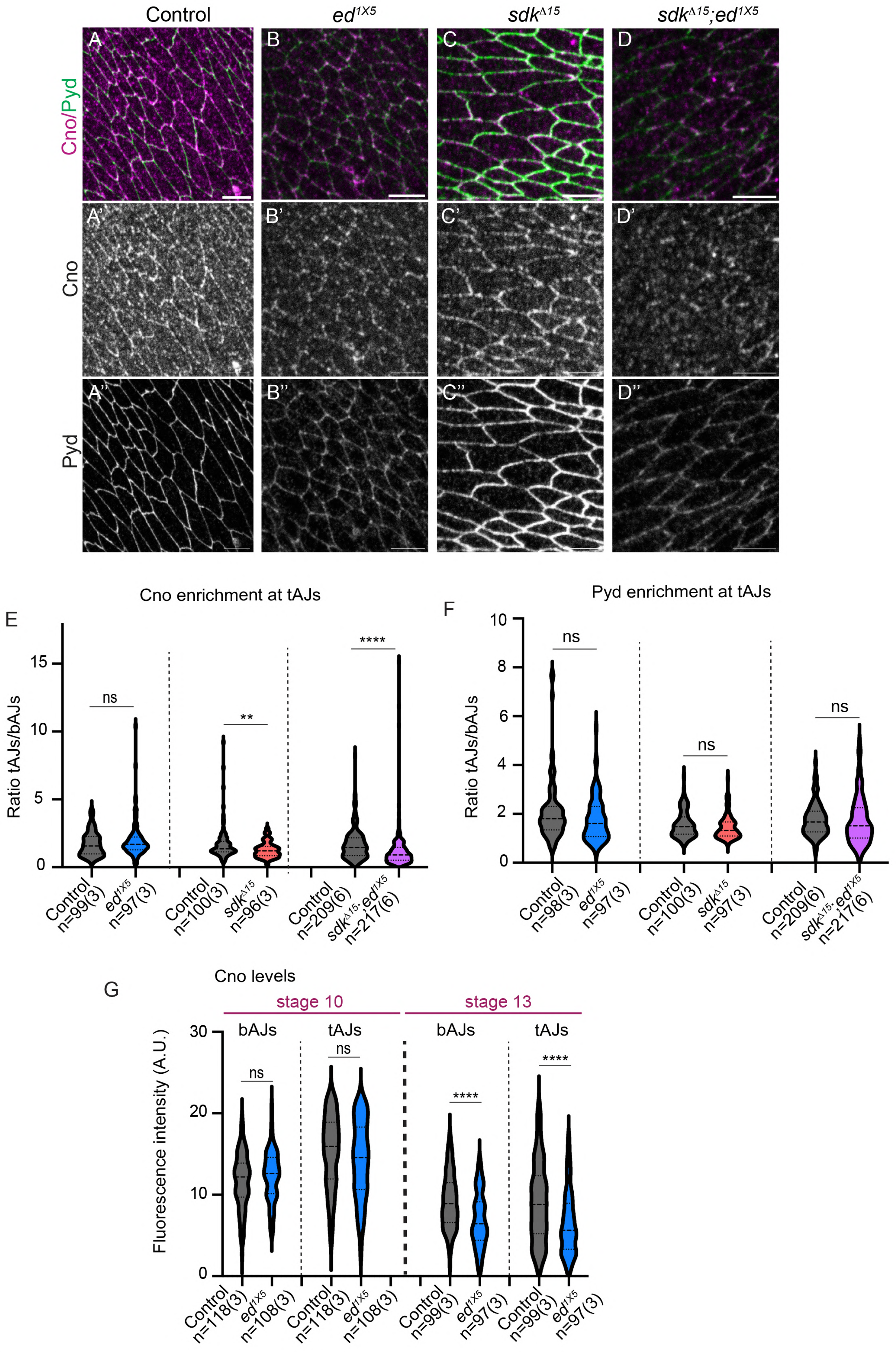
